# Elucidation of Amyloid-Beta’s Gambit in Oligomerization: Truncated Aβ fragments of residues Aβ1-23, Aβ1-24 and Aβ1-25 rapidly seed to form SDS-stable, LMW Aβ oligomers that impair synaptic plasticity

**DOI:** 10.1101/2022.12.04.519021

**Authors:** Beatriz Gil, Jamie Rose, Davide Demurtas, Gian-Filippo Mancini, Jessica Sordet-Dessimoz, Vincenzo Sorrentino, Nikita Rudinskiy, Matthew P. Frosch, Bradley T. Hyman, Marc Moniatte, Tara. L. Spires-Jones, Caroline E. Herron, Adrien W. Schmid

## Abstract

In Alzheimer’s disease (AD), Amyloid-beta (Aβ) oligomers are considered an appealing therapeutic- and diagnostic target. However, to date, the molecular mechanisms associated with the pathological accumulation or structure of Aβ oligomers remains an enigma to the scientific community. Here we demonstrate the strong seeding properties of unique Aβ fragment signatures and show that the truncated Aβ peptides of residues Aβ1-23, Aβ1-24 and Aβ1-25, rapidly seed to form small, SDS-PAGE stable assemblies of ∼5kDa to ∼14kDa molecular mass range. Mass spectrometry analysis of SDS-PAGE fractionated and gel extracted oligomers revealed that the truncated Aβ isoforms of residues 1-23 to 1-25 form stable entities with low molecular weight (LMW) oligomers, which strongly resemble the regularly reported Aβ entities of putative dimeric or trimeric assemblies found in human post-mortem AD and Tg mouse brain extracts. Furthermore, electrophysiological recordings in the mouse hippocampus indicate that LMW Aβ assemblies formed by fragments Aβ1-23 to Aβ1-25 significantly impair long-term-potentiation (LTP) in the absence of full-length Aβ1-42. Extensive antibody screening highlights the important observation, that the LMW Aβ assemblies formed by these truncated Aβ peptides escape immuno-detection using conventional, conformation specific antibodies but, more importantly, the clinical antibody aducanumab. Our novel findings suggest that there are new Aβ target “loopholes” which can be exploited for the development of therapeutic antibodies with binding properties against stable target hotspots present in Aβ oligomers. We provide here a first example of a new class of monoclonal antibody with unique binding properties against LMW Aβ oligomers, in the absence of binding to large fibrillar Aβ assemblies, or dense amyloid plaques. Our research supports a novel, unparalleled approach for targeting early, pathological Aβ species during the insidious phase of AD and prior to the appearance of large oligomeric or protofibrilar assemblies.

## Introduction

There is mounting evidence suggesting, that early therapeutic intervention will be the most promising therapeutic strategy in Alzheimer’s disease (AD) (1). As a consequence significant effort has been made in the past to gain insight into early, “molecular triggers” associated with pathological amyloid-beta (Aβ) aggregation, because Aβ oligomers are known to play a causative role in early synaptic dysfunction and cognitive decline in patients suffering from dementia (2). Therefore, Aβ oligomers are widely accepted as an appealing diagnostic and therapeutic target (3) in Alzheimer’s disease (AD). However, structural characterization of Aβ oligomers remains a challenging endeavour (4), which is mainly due to the labile and polymorphic nature found with these oligomeric entities. Consequently, it may not seem to be surprising, that the lack in clinical efficacy reported with earlier developed immunotherapies collectively reflect the intrinsic difficulties observed with the characterization of a “unique pathological target” associated with neurotoxic Aβ oligomers.

The aetiology and neurotoxic properties of different, low molecular weight (LMW) oligomeric Aβ species (4–9) such as Aβ dimers and trimers (10) or, larger oligomeric assemblies (11), have been subjected to thorough studies in the past, in order to identify the “holy grail” of Aβ assemblies as novel therapeutic targets in AD (12). Numerous studies have narrowed down the search of pathological Aβ species to LMW species, namely dimers (10,13,14) or trimers (11). More recently, McDonald and colleagues (15) reported that human AD brain extracts contain small Aβ structures of a 6–7 kDa molecular mass range, of which these Aβ species may form part of larger, neurotoxic Aβ assemblies. However, earlier attempts to identify specific Aβ isoforms associated with LWM Aβ assemblies and, more specifically, the ∼4kDa and ∼7kDa species, have failed (16).

Aβ peptide mediated neurotoxicity has been the subject of a host of *in vitro* and *in vivo* electrophysiological studies, indicating that acute exposure of both, Aβ42 (17) and the more soluble Aβ40 (18, 19) or, Aβ dimers isolated from human AD brains and CSF (14), significantly impair long-term-potentiation (LTP) in the rodent hippocampus. Earlier studies showed that intra-cerebral injection of Aβ40 in mice resulted in a rapid generation of truncated Aβ species (20), which resemble Aβ fragment fingerprints typically observed during *in vitro* Aβ oligomerization studies (12). The significance of truncated Aβ species is further supported by Mazzitelli and colleagues (21), who reported that the co-occurrence of the truncated fragment Aβ1-24 promotes Aβ42 aggregation and impairs Aβ peptide clearance in the mouse brain. In strong support with this concept, we provided strong analytical evidence on an aggregation induced auto-cleavage and hence accumulation of a specific Aβ fragment fingerprint, which can rapidly seed to form oligomeric Aβ assemblies (12). Taken together, these findings suggest that a gradual accumulation of different truncated Aβ isoforms, of which some bear C-terminal amidation (-CONH_2_), could play a central role in the early seeding events of Aβ oligomerization. These truncated Aβ species may therefore represent novel diagnostic markers as well as provide unprecedented therapeutic potential for the development of new antibody based AD therapies.

Using advanced technologies in quantitative and targeted mass spectrometry, together with our earlier developed neo-epitope antibody tools (12) against specific C-terminal truncated Aβ species, we were able to shed light on an enigma associated with LMW Aβ oligomers. Here we provide analytical proof of detection of specific truncated Aβ species present in LMW Aβ oligomers. This finding allows unprecedented and selective targeting of oligomeric assemblies, irrespective of oligomer structure or size. This gain in knowledge has led us to develop a novel, conformation selective mouse monoclonal antibody with unique binding properties against small oligomeric Aβ entities formed by the truncated Aβ isoforms Aβ1-23, Aβ1-24 and Aβ1-25, which appear during the early phase of Aβ peptide aggregation (12). In line with our antibody target design rational for early, LMW Aβ assemblies, we show that this novel antibody tool displays unique binding properties against LMW Aβ species, in the absence of binding to full-length Aβ42 monomers or higher order Aβ assemblies, such as amyloid plaques. We highlight here the strong need for identifying additional, molecular mechanisms associated with the early events of Aβ peptide aggregation during the long, insidious phase of AD. This in turn will allow the improvement of antibody-specific target engagement of early, pathological Aβ species present in the human AD brain and biofluids.

## Results

### Low molecular weight oligomers consist of an assembly of different C-terminally truncated Aβ isoforms

Numerous earlier reports show that oligomerization promotes the formation of labile Aβ assemblies that decompose into SDS-PAGE stable Aβ entities of dimeric, trimeric and tetrameric nature (12, 15). Here, we have used a combination of analytical tools to investigate the structural composition of these typical LMW Aβ assemblies observed during SDS-PAGE analysis. Preparation of Aβ42 oligomers was carried out following a standardized and widely applied protocol for the generation of amyloid derived diffusible ligands (ADDL) (22). In an initial approach, Aβ oligomers were analysed by SDS-PAGE followed by immuno-blotting, using the conventional antibodies 4G8 and 6E10, together with a selection of our earlier developed, highly selective neo-epitope antibody tools (Table I & Supplementary Data Fig. 1a&b.) against different C-terminally truncated Aβ species (12).

**Figure 1.**
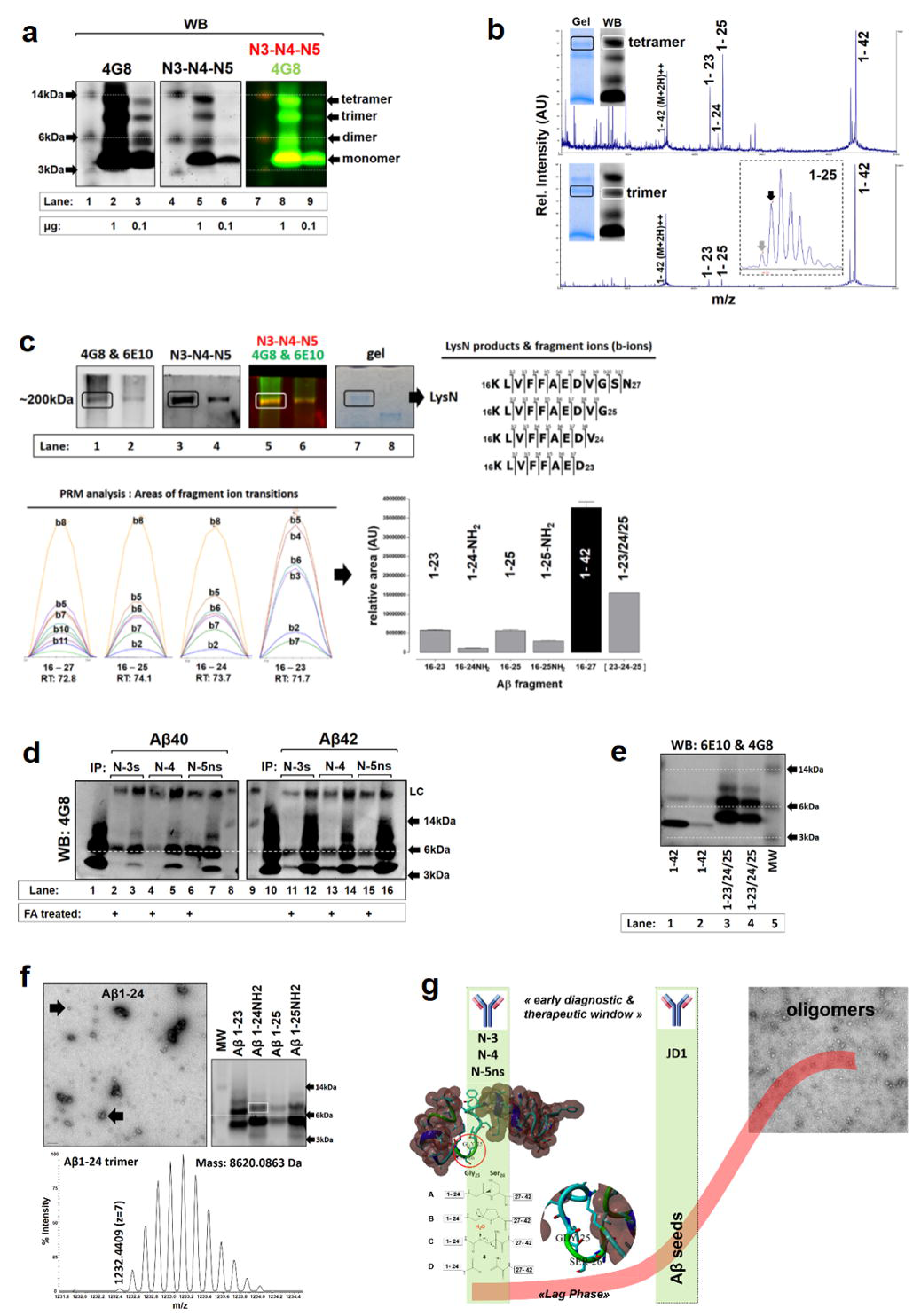
Truncated Aβ species rapidly seed to form small oligomeric entities of 6- 14kDa molecular weight range. **a** left; SDS-PAGE and immuno-blotting (4G8) analysis of Aβ42 ADDLs, showing the typical gel migration pattern of Aβ dimers, trimers and tetramers within the molecular mass range of 6-15kDa. Samples were loaded at high (1ug) and low (100ng) concentrations. Centre IB; re-probed membrane with a mix of neo-epitope antibodies (N3, N4 & N5ns) revealing the presence of C-terminal truncated fragments. (IB right: merged scan of both antibodies at different wavelengths: green=800nm (4G8), yellow=680nm (N3, N4 & N5ns). Neo-epitope antibodies bind to specific bands at the migration level of putative Aβ42 tetramers, trimers and dimers. Substantial neo-epitope antibody binding was also observed at the Aβ42 monomer level, which indicates the presence of C-terminally truncated Aβ species present in this band. **b** MALDI-TOF/TOF analysis of gel extracted (by diffusion) Aβ species found in tetramers (top) and trimers (bottom). Aβ1-25 was partially detected as a C-terminal amidated form (inset grey arrow), whereas Aβ1-24 was predominantly detected as the C-terminal amidated form (Aβ1-24-NH_2_). The strong MS signal intensities detected for Aβ23 and Aβ25, as compared to Aβ42 are in line with the IB signals found at the migration level of <14kDa (a). **c** WB and quantitative MS analysis of high molecular weight (HMW, ∼200kDa) Aβ assemblies detected in the insoluble fraction. WB’s were probed using either a combination of 6E10&4G (1^st^ blot, left) followed by re- probing with a mixture of neo-epitope antibodies (2^nd^ blot from left: N3, N4, N5ns & N5NH_2_) to detect the presence of fragments Aβ 1-23 to 1-25 in these HMW oligomers. Gel bands were excised at the migration level of HMW oligomers and were subjected to LysN digestion, giving rise to the mid region sequences (inset right) of either 16-27 (from full- length Aβ1-42 cleavage) or 16-25 (e.g. derived from Aβ1-25 cleavage product). LysN cleavage products were subjected to PRM analysis and the calculated total peptide areas, as a sum of all MS_2_ b-ion (b2 to b8) transitions (bottom, left), were used to estimate the relative abundance (histogram, right) of the different Aβ isoforms found within HMW oligomers. (Lanes 1-8 are identical Aβ samples but loaded at high (1, 3, 5) and low (2, 4, 6) concentrations, lane 8 = MWM). **d** WB analysis of Aβ40 (left) and Aβ42 (right) ADDLs. ADDLs were individually IP’ed with neo-epitope antibodies against fragments Aβ23, Aβ24 and Aβ25. Formic acid induced dissociation of ADDLs allowed a specific pull-down of these Aβ fragments (lanes: 2,4,6 and 11,13,15), whereas IP of native Aβ40 (lanes: 3,5,7) and Aβ42 (lanes: 12,14,16) ADDLs resulted in a pull-down of an oligomeric complex with substantial enrichment of a band centred at the migration level of ≥6kDa, which corresponds to a trimeric form of these Aβ fragments as seen by the migration pattern of a mixture of *in vitro* aggregated Aβ23 to Aβ25 (Figure 1e). (lanes: 1 & 10 = native Aβ40 & 42 ADDLs before IP, lanes: 8/9 = MW). **d** TEM imaging of the seeding properties observed with Aβ1-24, showing predominant spherical, oligomeric structures of 20-50nm diameter (arrows) and some clusters of larger aggregates (scale bar = 100nm). below; LC-MS analysis of Aβ24 trimers (WB inset) as seen by the multiply (m/z=1232.4409 M+H^7^) charged ion. Inset right; WB analysis of the gel migration behaviour of individually aggregated fragments of Aβ23, Aβ24 and Aβ25. All Aβ fragments form substantial amounts of trimers centred at the migration level of >6kDa (white box: Aβ24) as confirmed by LC-MS analysis. A shift in migration profile can be observed between dimeric and trimeric Aβ23 and Aβ24 assemblies, which accounts for the difference in molecular mass (200-300Da) found for the Aβ24 species (see table 2 for comparison). **e** WB analysis of Aβ assemblies formed by the mixture of fragments Aβ1-23, 1-24 and 1-25. The rapid seeding of these fragments results in the formation of a substantial population of dimers (≥5kDa) and trimers (≥6kDa), with some tetramers (≤12kDa). Monomers (≤3kDa) are no longer observed and the dimer (≥5kDa) band is slightly shifted when compared to the Aβ42 monomers at 4.5kDa, which accounts for the difference in molecular mass for the dimeric species. (lanes: 1/2 = Aβ42 and lanes: 3/4 = Aβ23, 24 & 25 mixture at high and low concentrations). **f** Summary of neo-epitope antibody properties and application (left green bar) for targeted analysis of early fragment fingerprints as a result of Aβ aggregation. The design and development rational for a new conformational specific antibody (JD1) (right green column) with binding specificity against LMW oligomers (early seeds) derived from early Aβ fragment fingerprints of Aβ1-23 to Aβ1-25 is shown.

**Table I.** Summary of the different antibodies used in the study. Antibodies no. 1 to5 are referred to neo-epitope specific antibodies in the study. IgM and IgG classes of the monoclonal antibody JD1 are shown, whereas the latter was mainly used in the study.

| No. | Antibody | Target / Epitope | Host | Type | Applications tested | Source |
| --- | --- | --- | --- | --- | --- | --- |
| 1 | N3 | Aβ C-term Asp23 | rabbit | polyclonal | IP, WB, DB | EPFL |
| 2 | N4 | Aβ C-term Val24-NH2 | rabbit | poyclonal | IP, WB, DB | EPFL |
| 3 | N5s | Aβ C-term Gly25-OH | rabbit | polyclonal | IP, WB, DB | EPFL |
| 4 | N5NH | Aβ C-term Gly25-NH2 | rabbit | polyclonal | IP, WB, DB | EPFL |
| 5 | N5ns | Aβ C-term 23-25 | rabbit | polyclonal | IP, WB, DB, ELISA, IHC | EPFL |
| 6 | JD1 | Aβ oligomer | mouse | monoclonal IgG1 | IP, DB, ELISA | EPFL |
| 7 | JD1 | Aβ oligomer | mouse | monoclonal IgM | DB, ELISA | EPFL |
| 8 | 6E10 | Aβ 3-8 | mouse | monoclonal | IP, WB, DB, ELISA, IHC | Covance |
| 9 | 4G8 | Aβ 17-24 | mouse | monoclonal | IP, WB, DB | Covance |
| 10 | MM26-2 | Aβ C-term 42 | mouse | monoclonal | WB | Anaspec |
| 11 | A11 | oligomers | rabbit | polyclonal | DB | Thermo-Fisher |
| 12 | OC | Aβ aggregates | rabbit | polyclonal | DB, IHC | Merck |
| 13 | Aducanumab | Aβ oligomers / fibrils | recombinant | human IgG | DB | Creative Biolabs |
| 14 | Neurogranin | - | sheep | polyclonal | WB | R&D Systems |
| 15 | Somatostatin | - | rabbit | polyclonal | WB | Abbexa |
| 16 | Syn-1 | alpha-synuclein | mouse | monoclonal | WB | BD Tansduction Labs |
| 17 | PSD94 | - | mouse | monoclonal | WB | abcam |
| 18 | b-actin | C-term | mouse | monoclonal | WB | Santa Cruz Biotech. |
| 19 | homer 1 | - | rabbit | polyclonal | WB | Synaptic Systems |

In line with earlier studies we show that, under denaturing SDS-PAGE conditions, large oligomeric assemblies dissociate into LMW species of putative dimeric, trimeric and tetrameric Aβ species (Fig. 1a). Probing the WB membrane with a combination of neo- epitope antibodies N-3, N-4 and N-5ns against Aβ fragments of residues 1-23 (mw: 2775Da), 1-24 (mw: 2875Da) and 1-25 (mw: 2932Da) respectively, allowed identification of C-terminally truncated Aβ species at the migration level corresponding to Aβ42 monomers, dimers, trimers and tetramers. Detection of these truncated Aβ fragments appeared to be significantly increased in Aβ42 tetramers when compared to trimers and dimers, whereas strong antibody signals were also observed at the Aβ42 monomer level (mw: 4512Da).

To confirm the identification of different Aβ species in LMW oligomers, we excised SDS-PAGE gel bands at the migration level of Aβ42 monomers to tetramers (Fig. 1b, inset: gel), followed by a repeated gel dehydration & rehydration process to allow diffusion of the gel trapped Aβ species into the solution. Following gel extraction, the solution containing different Aβ species was cleared and concentrated using C18 stage tips and subjected to MALDI-TOF/TOF MS analysis. MS analysis revealed a high abundance of fragments Aβ1-23 and Aβ1-25 together with full-length Aβ42 in the tetramer gel band (Fig. 1b; top spectrum). Similarly, the C-terminally amidated (-NH_2_) forms of fragments Aβ1-24 and Aβ1-25 were also identified, however at significantly lower abundance. Overall, the MS signal intensities found for Aβ1-23 and Aβ1-25 were in line with the immuno-blot signal observed at the identical gel migration level of Aβ42 tetramers (≤15kDa) (Fig. 1a, lanes: 5&8). We could also identify similar Aβ fragment signatures in the trimer gel band, however, at significantly lower ratios when compared to tetramers, which corresponds to the decreased immuno-blotting signals found at the corresponding migration level.

In addition to the above outlined diffusion extraction procedure, an identical copy of the gel bands was subjected to in-gel digestion, using LysN protease followed by LC-MS analysis, to further corroborate the findings by WB and MALDI-TOF MS, as well as to allow identification of additional Aβ isoforms present within these SDS-PAGE-stable LMW oligomers. High-resolution LC-MS analysis revealed the presence of a diversity of N-terminally, as well as C-terminally modified Aβ species across the different gel migration levels (Supplementary Data Fig. 1c-e). Generally, identification of fragments Aβ1-23, Aβ1-24 and Aβ1-25 could be confirmed in the monomer, dimer, trimer and tetramer bands and these fragments were found to be significantly enriched in the monomer and tetramer bands respectively, which is in line with the WB and MS analysis shown above (Fig. 1a). The increased abundance of truncated Aβ species detected at the migration level of 3-5kDa may result from a gel migration induced dissociation of Aβ assemblies, giving rise to the monomeric and dimeric form of Aβ1-23, Aβ1-24 and Aβ1-25 (Table II). Moreover, LC-MS analysis allowed for unbiased identification of the C-terminally amidated fragment Aβ1-24 (Val_24_). In addition to the above listed truncated forms, we were also able to identify C-terminal amidation at positions His_13_, His_14_ and Asp_23_. Modifications associated with the far C-terminal consisted of a repertoire of specific truncations found at position Gly_37_, Gly_38_, Val_39_ as well as Ile_41_ (Supplementary Data Fig. 1e).

**Table II.** List of Aβ peptide fragments used in the study. Theoretical masses of Aβ dimers, trimers and tetramers was calculated using the average peptide mass.

| Aβ Peptide | C-term. | Abbreviation | monoisotopic mass (Da) | average mass (Da) | Dimer (Da) | Trimer (Da) | Tetramer (Da) |
| --- | --- | --- | --- | --- | --- | --- | --- |
| 1-23 | COOH | 23 | 2775,27 | 2776,96 | 5554 | 8331 | 11108 |
| 1-24-NH2 | CONH2 | 24N | 2873,35 | 2875,11 | 5750 | 8625 | 11500 |
| 1-25 | COOH | 25 | 2931,36 | 2933,15 | 5866 | 8799 | 11733 |
| 1-25-NH2 | CONH2 | 25N | 2930,37 | 2932,16 | 5864 | 8796 | 11729 |
| 1-40 | COOH | 40 | 4327,15 | 4329,86 | 8660 | 12990 | 17319 |
| 1-42 | COOH | 42 | 4511,27 | 4514,10 | 9028 | 13542 | 18056 |

### Truncated Aβ species may seed to form SDS-PAGE stable entities with large Aβ oligomeric assemblies of a 50-200kDa molecular mass range

We reasoned that the detection of small Aβ assemblies at the migration level of 5-14kDa may result from the dissociation of larger (≥30kDa), metastable Aβ entities. We therefore decided to analyse the insoluble fraction after centrifugation of the ADDLs (soluble fraction) preparation. This insoluble fraction is known to contain mainly large Aβ assemblies of protofibrillar and fibrillar nature (data not shown). WB analysis of this fraction revealed the presence of different, large Aβ assemblies at the migration level of 50kDa to 200kDa (Supplementary Data Fig. 2a). Probing the membrane with antibodies directed against the N-terminal (6E10) and mid-region (4G8) or, the C-terminal ending of Aβ42 (MM26-2) (Table I) allowed the identification of the bulk of Aβ assemblies centred at the LMW migration level of 4-14kDa, as observed with the soluble ADDL fraction shown above. However, application of a cocktail of neo-epitope antibodies (N3, N4, N5ns and N5NH2) also strongly indicated the presence of distinct Aβ assemblies found at the gel migration level of ∼200kDa (Fig. 1c) as well as ∼98kD, ≥62kDa and ≥50kDa (Supplementary Data Fig. 2a). We observed particularly strong binding of 6E10 & 4G8 as well as N3-N5 (mix) to a large Aβ entity found at ∼200kDa (Fig. 1c, top: WB). We therefore decided to carry out in-gel digestion followed by quantitative MS analysis of these large Aβ entities in order to confirm the presence of the different Aβ isoforms of Aβ1-23 to Aβ1-25 within these stable assemblies. Our quantitative MS analysis confirmed the high abundance of the different Aβ isoforms of 1-23, 1-24NH_2_, 1- 25 and 1-25NH_2_ present in these large Aβ assemblies (e.g. 200kDa) of which the overall calculated sum of all truncated Aβ isoforms accounted ≥40%, when compared to full-length Aβ1-42 (Fig. 1c, bottom: histogram). We also provide quantitative MS data on the different Aβ isoforms detected at the gel migration level of ≥50kDa, ∼98kDa and ∼200kDa (Supplementary Data Fig. 2b).

**Figure 2.**
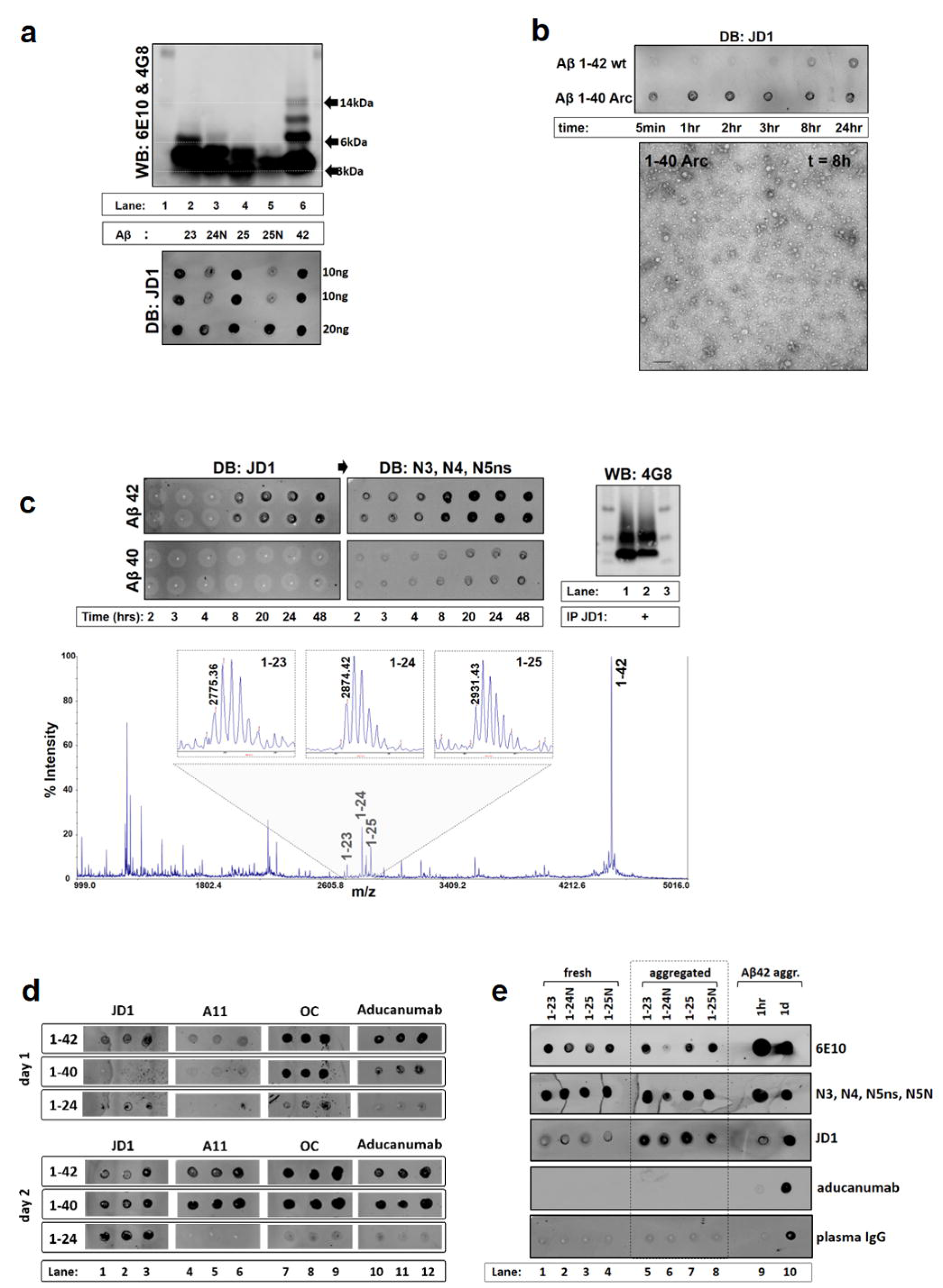
LMW Aβ oligomers escape immuno-detection using conventional, conformation specific antibodies. **a** WB (4G8 & 6E10) and DB (JD1) analysis of the migration profile of JD1 positive species formed by aggregated Aβ1-23 to Aβ1-25 (lanes: 2- 4) and full-length Aβ42 (lane: 6). The presence of Aβ42 (lane 6) species at the migration level of dimers and trimers is revealed by WB analysis. Below; DB validation of JD1 positive species. JD1 positive signals found for Aβ42 were associated with the presence of bands detected at ≥6kDa and ≤14kDa (lane: 5). **b** Aggregation kinetic analysis of the mutant Aβ40- Arc versus wt Aβ42 peptide using JD1 (DB) and TEM imaging of Aβ40-Arc oligomers at 8hrs. Aβ entities of spherical (10-50nm) and small protofibrillar morphology can be observed with Aβ40-Arc (scale bar = 100nm). **c** Monitoring of Aβ42 and Aβ40 aggregation kinetics using JD1 and MS. top: DB validation using JD1 (left) and re-probed with a mixture neo- epitope antibodies (right): positive JD1 signals are observed following 8hrs of Aβ42 aggregation (DB) and this was accompanied by an appearance of fragments Aβ23, Aβ24 and Aβ25 as probed with N3, N4 and N5ns (right). Bottom: MS (MALDI-TOF-TOF) analysis of Aβ fragment signatures detected at t=8hrs. Inset right; WB analysis of IP’ed (JD1) Aβ42 (t=8hrs) (lane 2) showing the pull-down of a predominant band centred at the migration level of ≥6kDa. (lane 1: Aβ42 starting material). **d** DB validation of Aβ40 and Aβ42 as well as truncated Aβ1-24 aggregation kinetics using JD1 (Lanes: 1-3), the oligomer specific antibody A11 (lanes: 4-6), conformation specific antibody OC (lanes: 7-9) and Aducanumab (lanes: 10-12) (samples were spotted at two different concentrations: lanes 1-2, 4-5, 7-8,10-11 = 50ng and lane 3, 6, 9, 12 = 100ng). A clear difference in antibody binding to aggregated Aβ24 can be seen between JD1 and A11 as well as Aducanumab. **e** Validation of JD1, Aducanumab as well as human plasma, natural IgGs or 6E10 binding specificity against freshly prepared (lanes: 1-4) or aggregated (lanes: 5-8) forms of Aβ fragments Aβ23, Aβ24 and Aβ25 as well as Aβ42 (lanes: 9-10). JD1 has strong binding affinity for the aggregated form of the different Aβ fragments; whereas no binding was observed with Aducanumab or plasma IgGs (lanes. 5- 8) against the soluble or aggregated form of these C-term truncated Aβ species.

### Dissociation of Aβ oligomers results in the isolation of Aβ species of 6kDa to 14kDa mass range

Given the observation that truncated Aβ species form stable complexes with LMW and HMW oligomers, we were interested in validating if we are able to pull-down Aβ oligomers formed by Aβ40 or Aβ42 peptides. To address this question, we prepared ADDLs of which one part of the preparation was subjected to 70% formic acid (FA) denaturing (3hrs at room temp.) followed by IP using neo-epitope antibodies N-3s, N-4 or N-5ns (Table I & Supplementary Data Fig. 1a&b). We show that selective IP of FA dissociated ADDLs resulted in a specific capturing of two Aβ species centred at the gel migration level of 6kDa and <14kDa, whereas IP of native (not denatured) ADDLs resulted in the pull-down of the typical oligomeric assemblies found at a migration level of 6-14kDa (Fig. 1d). The selective pull-down and therefore isolation of the ≥6kDa Aβ species was found to be more straightforward with Aβ40 assemblies (Fig. 1d, left; lanes 2-7), whereas Aβ42 ADDLs appeared to be more resistant to FA dissociation (Fig. 1d, right; lanes: 11-16).

### Aβ fragments of Aβ1-23, Aβ1-24 and Aβ1-25 rapidly seed to form SDS-stable, LMW assemblies in the absence of full-length Aβ1-42

Because our earlier studies strongly indicated that Aβ aggregation is accompanied by the appearance of the typical fragment fingerprint consisting of truncations at residues 1-23, 1-24 and 1-25, we were interested in understanding the seeding properties using a mixture of these C-terminally truncated Aβ species. We show that, *in vitro* aggregation of these truncated Aβ fragments results in the rapid formation of stable assemblies within the 5-15kDa molecular range, of which dimers (≥5kDa) and trimers (≥6kDa) are the most predominant species formed by these Aβ fragments, whereas monomers (theoretical av. mass range: 2776Da to 2933kDa) were no longer detected (Fig. 1e, lanes: 3-4). Consistent with this observation, earlier reports showed a rapid seeding behaviour with fragment Aβ1-28, revealing gel migration profiles that are consistent with stable Aβ1-28 dimers (22). The gel migration profile of the peptide mixture of Aβ1-23 to Aβ1-25 dimers (>5kDa) was significantly shifted when compared to full-length Aβ1-42 monomers (4.5kDa) (Fig. 1e, lanes 1-2), whereas the trimeric form of these truncated Aβ species (theoretical mass: 8-9kDa) was found at a migration level above ≥6kDa, which strongly resembles the migration profile of earlier reported putative Aβ42 dimers (10) or 4-7kDa assemblies (15) isolated from human AD brain extracts.

We then investigated the self-seeding properties of individual truncated Aβ species Aβ1-23, Aβ1-24-NH_2_ and Aβ1-25 as well as the C-term amidated form of Aβ1-25 (NH_2_). In vitro aggregation of Aβ1-24 showed that this fragment has rapid self-seeding properties, resulting in the formation of SDS-PAGE stable assemblies in the MW range of dimers to trimers (Fig. 1f, WB inset). Transmission electron-microscopy (TEM) imaging revealed that fragment Aβ1-24 preferentially seeds to form homogenous spherical aggregates with some clusters of small protofibrillar structures. The gel migration profile of Aβ1-24 and Aβ1-25 peptides were similar (Fig. 1f), whereas the gel migration profile appeared to be slightly shifted when compared to fragment Aβ1-23, which may account for the difference in MW between these Aβ species (Aβ23 vs. Aβ24 dimer = 198Da, Aβ24 dimer vs. Aβ25 dimer = 114Da) (Table II). The true molecular entity of these small oligomeric assemblies observed by SDS-PAGE was further confirmed using high-resolution LC-MS analysis, revealing the presence of highly stable Aβ1-24 trimers (8.6kDa) (Fig. 1f, bottom: MS spectrum) as well as dimers (5.7kDa) or even pentamers (14.3kDa) (Supplementary Data Fig. 2c-d). The MS analysis of stable dimers to pentamers of these truncated Aβ peptides further corroborates the migration pattern observed by denaturing SDS-PAGE analysis (Fig.1f, WB inset), and therefore rules out any analytical artefacts associated with gel migration (23).

### Detection of early, pathophysiological Aβ seeds requires the development of novel antibody tools, with high binding selectivity for small oligomeric assemblies, in the absence of monomer binding

The above reported findings clearly highlight the need for the development of additional antibody molecules with high binding specificity for LMW Aβ oligomers, irrespective of the presence of the full-length Aβ42 or Aβ40 target. To address this limitation, we set out to develop a conformational specific antibody; with preferential binding properties against a repertoire of LMW Aβ assemblies formed by here described truncated Aβ species. We speculated that the application of such a conformation specific antibody would serve as an additional analytical tool as compared to our earlier developed neo-epitope antibodies, therefore allowing the detection of the gradual appearance of small Aβ entities, prior to the accumulation of large, metastable oligomeric or protofibrillar species (Fig. 1g).

We therefore immunized mice using a synthetic, oligomeric form of fragment Aβ1-24 and screened clones for antibodies with binding properties against a linear (denatured), and aggregated (positive target) form of the different C-terminally truncated Aβ fragments of interest as well as against denatured, full-length Aβ42 and Aβ40 peptide (negative target). Using a combination of ELISA and DB techniques, we were able to identify six antibody-secreting clones (data not provided), with different antibody binding properties against oligomeric Aβ assemblies of interest, of which the properties of one selected monoclonal antibody (mAB), referred to as **“*JD1*”** is described below.

### The monoclonal antibody JD1 binds LMW Aβ assemblies, which appear during the early time course of Aβ aggregation

To characterize the binding specificity of JD1, we initially compared JD1 binding properties with the two conventional antibodies 4G8 and 6E10, as well as different, conformation specific antibodies such as the A11 or OC antibody. We found that the intrinsic binding property of JD1 is not limited to fragment Aβ1-24, since JD1 binding was also observed for the different aggregated forms of fragments of Aβ1-23 and Aβ1-25-NH_2_ (Fig. 2a; bottom DB). Immuno-blotting analysis indicated that JD1 positive signals found by dot blotting are in agreement with the presence of LMW oligomeric assemblies found at the Aβ peptide migration range of dimers and trimers (5-14kDa) and this could be also confirmed using aggregated Aβ42 (Fig. 2a, top WB, lane: 6). While JD1 clearly binds to oligomeric Aβ assemblies using native DB, we do not observe positive JD1 binding of these LMW species by WB. The reason for this observation may result from either dissociation of larger (≥ 14kDa) Aβ assemblies or from a SDS-PAGE induced denaturing of the conformational epitope.

DB analysis of Aβ aggregation kinetics indicated that significant JD1 binding can be observed following ≥8hrs aggregation of the full-length, wild type (wt) Aβ1-42 peptide (Fig. 2b). Remarkably, aggregation kinetics analysis of the arctic mutant Aβ40 revealed that, JD1 binding significantly preceded the detection of oligomeric assemblies formed by Aβ40-Arc peptide, when compared to wt Aβ42 peptide. TEM imaging indicated that selective JD1 binding was accompanied by the presence of predominantly oligomeric entities found with Aβ40-Arc (Fig. 2b), which is in line with the morphology observed with fragment Aβ1-24 (Fig. 1f). Probing Aβ42 peptide aggregation with neo-epitope antibodies revealed that the rise in JD1 positive species is generally associated with a gradual accumulation of C-terminally truncated Aβ species of residues 1-23 to 1-25 and this was further confirmed by MALDI-TOF MS (Fig. 2c, bottom: MS spectrum), whereas no antibody binding was observed for the more soluble, wt Aβ40 peptide during the same time course of peptide aggregation (Fig. 2c; top DB).

### Validation of JD1 binding properties against oligomeric Aβ species and its potential applications for screening methods

We show that detection of JD1 positive species was accompanied by the gradual accumulation of a typical Aβ fragment fingerprint (Aβ23 to Aβ25) as shown by the application of neo-epitope antibodies (Supplementary Data Fig. 3a) and JD1 proved to be useful for screening aggregation inhibitors, such as the green tea extract (EGCG) (Supplementary Data Fig. 3b). In line with the DB screening we show that, IP-MS analysis with JD1 does not result in the pull-down of soluble, Aβ42 monomers at early time points (t ≤ 30min), when compared to prolonged aggregation times (t = ≥2hrs) (Supplementary Data Fig. 3c). We also took advantage of the C. elegans GMC101 worm model to further understand the binding property of JD1 using a living organism. GMC101 worms constantly express the human Aβ42 isoform in muscle cells and develop age-progressive paralysis and significant amyloid deposition which are A11 positive species (24). WB analysis of aged GMC101 worms revealed a significant accumulation of different LMW Aβ species at the molecular weight range of 5-14kDa. IP-MS analysis of GMC101 extracts confirmed the abundance of different C-terminal truncated Aβ species (Supplementary Data Fig. 3d), which reflects the typical Aβ fragment signatures found in Tg2576 mice and human post-mortem brain tissue extracts (25). Furthermore, we show that JD1 also binds oligomeric assemblies formed by synthetic mouse Aβ42 sequence, though, with significantly lower affinity (Supplementary Data Fig. 3e), whereas, no binding was observed against the reversed human Aβ42-1 sequence (Supplementary Data Fig. 3f) or aggregated alpha-synuclein protein (data not shown). These observations collectively rule out JD1’s binding property against oligomeric assemblies formed by other amyloidogenic proteins, as observed with earlier developed conformation specific antibodies (26). Collectively these observations highlight JD1’s unique binding property for predominantly, small oligomeric Aβ assemblies, in the absence of binding of full-length Aβ1-42 or Aβ1-40 monomers.

**Figure 3.**
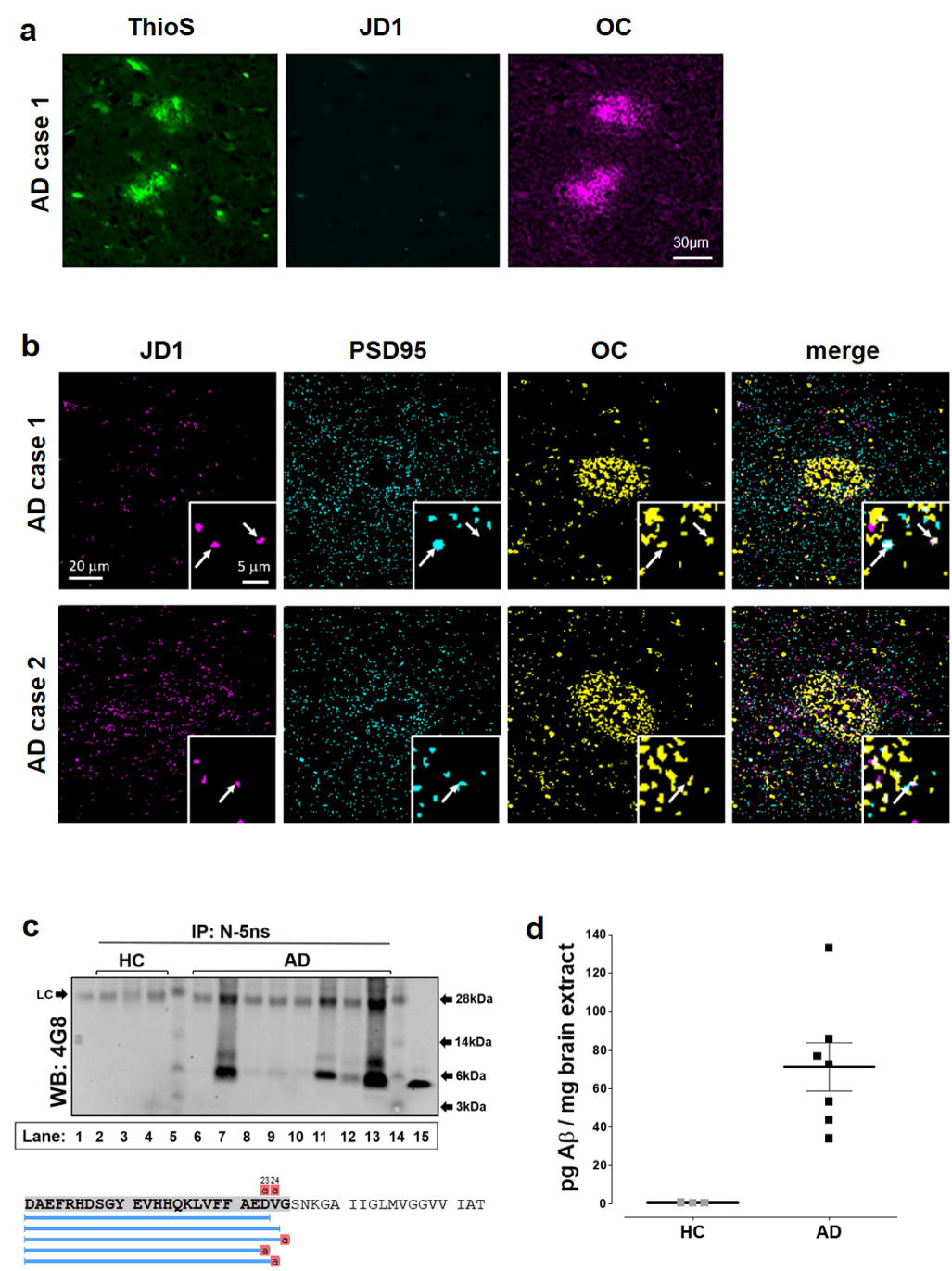
JD1 positive oligomers are detected in the post-synaptic vicinity of AD brains. **a** IF staining of amyloid plaques found in a human AD brain, using Thioflavin S (Thio S), JD1 and OC antibody. Significant plaque staining was observed with Thio S, which co- localized with OC staining, whereas no staining of dense core plaques was found using JD1. **b** Array tomography images of Aβ deposits detected in two post-mortem human AD brains using JD1, anti-PSD95 and OC antibody (left to right). JD1 positive species were detected in the post-synaptic vicinity, which co-localize with PSD95 positive staining. Large images are maximum intensity projections of 5 serial sections and insets are single sections confirming colocalization. (for brain tissue details see Supple. Table I). **c** Example of an IP-WB analysis of human cortical brain tissue extracts using the neo-epitope antibody N5ns. Significant pull-down of dimeric and trimeric Aβ species are observed in a majority of AD (lanes 6-14) but not control (lanes 2-4) brains. (lane 1: IgG ctrl, lane 15: MW, Lane 16: aggregated synthetic Aβ25 dimer). Below; Representative summary of an IP-MS analysis of an AD brain sample (lane 14) using a mixture of antibodies N3, N4 and N5ns. Blue bars highlight the sequences of all Aβ species identified (FDR ≤1%) (PEAKS Studio soft.) **d** ELISA analysis of JD1 positive assemblies found in post-mortem human AD (n=7) brains (FA extracts) but not in age matched controls (HC, n=3). (range: 35pg-132pg / mg brain tissue) (median ± SEM of technical triplicates).

### Small Aβ assemblies formed by the N-terminal fragments Aβ1-23 to Aβ1-25 are not detected by conventional, anti-oligomeric antibodies or natural plasma IgG’s

Due to the strong seeding properties observed with C-terminally truncated Aβ species, we investigated if these LMW oligomeric assemblies can be detected using conventional, anti-oligomeric antibody tools. For this purpose, full-length Aβ42 and Aβ40, as well as the C-terminally amidated fragment Aβ1-24-NH2 were subjected to aggregation kinetics and probed with the “anti-oligomer” specific antibody A11 and the “conformation-specific” antibody OC (6,26,27), as well as a biosimilar of the clinical antibody aducanumab (28, 29). No visible antibody binding was observed at day 0 (data not shown), which is in line with earlier reported antibody screening studies using A11 and OC (30). Following one day of Aβ peptide aggregation, significant antibody binding was only observed against peptides Aβ42 and Aβ40 using the OC and aducanumab antibodies (Fig. 2d), whereas A11 reached a similar level of binding significance following two days of aggregation. Generally, positive immuno-probing signals for Aβ42 preceded the detection of Aβ40 oligomers, which is likely due to the increased aggregation propensity found with Aβ42. At day two, JD1 binding to aggregated Aβ42 and Aβ40 was similar when compared to aducanumab, but with a noticeable lag in detection of Aβ40 at day 1. Intriguingly, our screening studies revealed a striking difference in antibody binding affinity for the aggregated Aβ1-24 peptide. Detection of oligomeric Aβ1- 24 assemblies was significantly reduced using the OC or aducanumab antibody and absent with A11 (Fig. 2d & Supplementary Data Fig. 3g). To determine if the intrinsic binding properties of JD1 are associated with an oligomeric or linear, denatured form of Aβ peptide, we prepared fresh Aβ peptide standards by dissolving and sonicating the Aβ peptides in HFIP solution, followed by solution evaporation and solubilization in a 50mM NaOH solution as described earlier (30). Sample preparation of freshly prepared (t=0) or aggregated (t=1d) Aβ peptide standards where screened by DB using JD1 and aducanumab, together with a cocktail of neo-epitope antibodies (N3, N4, N5ns, N5N) as well as 6E10 for the detection of all Aβ assemblies. Our findings strongly indicate that JD1 has significantly increased binding properties for the aggregated form of fragments Aβ1-23, Aβ1-24-NH_2_, Aβ1-25, Aβ1-25NH_2_ and that detection of these Aβ species was significantly reduced with aducanumab (Fig. 2e, lanes: 5-8). Application of neo-epitope antibodies enabled the detection of both, denatured as well as aggregated forms of the Aβ peptide, indicating that these antibodies bear high C-terminal binding specificity, irrespective of Aβ peptide structure, which is in agreement with our earlier reported validation studies (12).

Human plasma is known to contain natural IgGs antibodies that are reactive against a broad range of different amyloidogenic species. Because natural antibodies have been reported to bear neuroprotective properties against Aβ peptide (31–33), we were interested in understanding if these plasma IgG’s also bear binding properties against the here described LMW Aβ oligomers. In-spite of the high, natural abundance of conformation-specific antibodies reported in human plasma, we could not identify plasma IgG selective binding to the small oligomeric Aβ entities of fragments Aβ1-23 to Aβ1-25. However, significant plasma IgG binding was observed for aggregated, but not soluble Aβ42 peptide and levels of plasma IgG binding was comparable with aducanumab (Fig. 2e, lane: 10). Overall, these observations highlight the important finding that, the small oligomeric Aβ assemblies formed by the truncated fragments Aβ1-23 to Aβ1-25, may escape conventional immuno-based detection, using the oligomer-specific antibody A11 or aducanumab.

### Identification of JD1 positive assemblies in human post-mortem AD brains highlights the pathological significance of LMW Aβ assemblies in AD

To further provide evidence of the significance of LMW oligomeric Aβ species in AD brains, we decided to screen biological samples using a combination of immuno-fluorescence (IF) and array tomography imaging as well as IP-MS and ELISA assays. Having confirmed the presence of truncated Aβ fingerprints in a set of human brain samples earlier (12), we set out in specifically identifying JD1 positive species in post-mortem human brains.

IF imaging of human AD brains revealed the presence of dense core plaques, which were thioflavin S (ThioS) and OC positive, however, we did not observe any JD1 positive staining of these dense amyloid plaques (Fig. 3a). Therefore, we were interested in identifying the presence of JD1 positive species at synapses, because one of the early and important effects of oligomeric Aβ in AD, is the binding of oligomeric Aβ to synapses, resulting in synaptic dysfunction and gradual loss of synapses. Tissue from human subjects was prepared for high-resolution ribbon array tomography as reported previously (34), allowing accurate and high-resolution detection of individual synapses. Similar to our IF analysis, application of the OC antibody resulted in strong labelling of dense core Aβ plaques as well as individual postsynaptic punctae, which co-localized with PSD95 protein staining. No JD1 positive staining could be observed for dense Aβ plaques, which is in line with the IF analysis of aged (≥9months) Tg2576 mouse brains (data not shown). Interestingly, we clearly observed selective JD1staining at individual postsynaptic punctae, which co-localized with OC and PSD95 staining (Fig. 3b). This observation indicates that JD1 positive Aβ assemblies may represent early, neurotoxic Aβ species associated with synapse binding and therefore resulting in gradual neurodegeneration.

### Targeted mass spectrometry analysis of human AD brain extracts reveals the presence of LMW Aβ assemblies, consistent with the detection of fragments Aβ1-23 to Aβ1-25

We decided to analyse human post-mortem brains to further support the results of our high-resolution array-tomography and provide insight into the pathophysiological significance of these truncated Aβ isoforms as well as the LMW oligomers. We employed IP-WB and IP-MS analysis of eight AD and three human control (HC) brain extracts (Supplementary Data Table I) and found that IP with N-5ns enabled the isolation of Aβ species centred at a gel migration level of 5-6kDa and ≥6kDa, which corresponds to our above observed dimeric and trimeric species of fragment Aβ1-25 (Fig. 1e). Significant levels of dimeric and trimeric Aβ species were detected in four AD brains (Fig. 3c, lanes: 7, 12 13 & 14), whereas the detection of these Aβ species was significantly lower in three other AD subjects (Fig. 3c, lanes: 8-10). No Aβ was detected in brains from human controls (Fig. 3c, lanes: 2-4) as well as in one AD subject (lane: 6). Generally, IP-WB analysis of these AD brain extracts strongly resembled the migration profile of the above reported FA dissociated ADDLs (Fig. 1d). Moreover, using IP-MS analysis with a neo-epitope antibody cocktail (N3, N4, N5ns), we were able to confirm the overall presence of different Aβ species in AD brain extracts but not human controls (Fig. 4c; bottom inset & Supplementary Data Fig.4a-e). Overall, our IP-MS analysis allowed the unbiased identification of different truncated Aβ species of residues 1-23, 1-24, 1-25, of which fragment 1-24 was predominantly detected as a C-terminal amidated (-CONH_2_) form. In addition, we were also able to identify the C-terminally amidated fragment of Aβ1-25 (Supplementary Data Fig. 4e) using a recently developed neo-epitope antibody (N-5NH2) (Table I).

**Figure 4.**
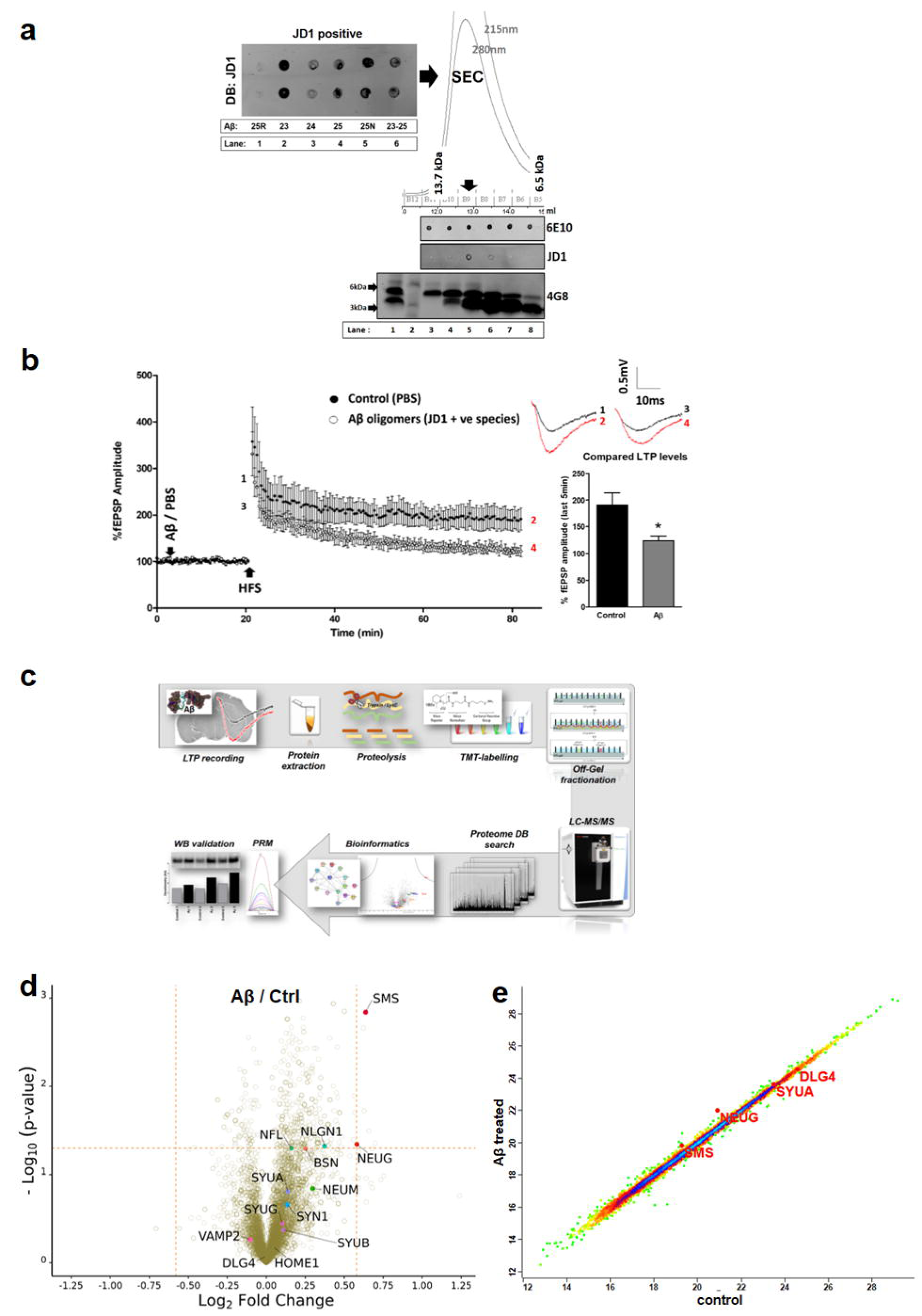
LMW oligomeric assemblies derived from C-terminally truncated Aβ species significantly inhibit LTP in the mouse hippocampus. **a** left; representative DB screening of JD1 positive assemblies formed by the individual fragments Aβ23 to Aβ25 (lanes 2-5) as well as using a mixture of all fragments (lane 6). A reverse sequence of Aβ25-1 was used to show that aggregation of this sequence does not generate JD1 positive assemblies (lane 1). Right; SEC (Superdex 75, 10/300) and DB analysis of LMW assemblies formed by the mixture of Aβ fragments 1-23 to 1-25. JD1 positive assemblies were detected in SEC fraction volume 13ml, which contained dimeric and trimeric (≥6kDa) assemblies of these fragments (WB inset below), whereas monomers (≤3kDa) eluted in fraction volume 16ml. **b** Effect of application of LMW Aβ oligomers (SEC purified JD1 positive assemblies) on LTP in hippocampal slices from young wild type mice. Acute application of Aβ oligomers (500nM) to hippocampal slices significantly depressed LTP amplitude compared to control conditions (P<0.05). (inset right: top) Sample traces represent fEPSP before (**1**) and 60min after HFS (**2**) in control LTP, and before (**3**) and after HFS (**4**) Aβ-mediated LTP. (inset right: bottom) Bar chart comparing magnitude of LTP in control (190.6 ±22.94) and Aβ conditions (123.8 ±9.05) (N=5). All data are presented as Mean ±SEM and analyzed using unpaired Student’s t-test. JD1 positive Aβ oligomer treated sliced (open circles) as compared to PBS controls (filled black circles). **c** Diagram of the analytical workflow applied for mouse hippocampal tissue extraction post-LTP recordings, protein digestion, isobaric labelling (TMT), off-gel isoelectric focusing, MS analysis and bioinformatics as well as orthogonal hit validation by targeted, quantitative MS and immuno-blotting. **d** Volcano plot analysis (y-axis of volcano plots: -Log10 p-value from two-sided T test corrected by Benjamini-Hochberg procedure) of differentially regulated proteins (x-axis: Log2 fold-change) in mouse brain tissue (n=3), following treatment with Aβ oligomers versus PBS controls. The dashed lines indicate a threshold of ± Log2 0.58 fold change (vertical lines) and a 0.05 p-value cut-off threshold (horizontal line). At the given cut-off, somatostatin (SMS) appears to be significantly up- regulated (FC= Log2 0.63) in Aβ treated brains. A similar trend in FC was observed for neurogranin (NEUG), however at significantly reduced level of significance (FC= log2 0.58). Proteins were graphed by fold change (FC, difference) and significance (-Log p) using a false discovery rate of 0.05 and an S0 of 0.1. Protein ID’s in red (SMS and NEUG) were identified as significantly up-regulated (ggVolcanoR tool), whereas only SMS passed the confidence level of P<0.05 using Perseus software (Supplementary Data Figure 9). **e** Representative example of a regression analysis of (intra-TMT channel) protein correlation (mouse #2) for control versus Aβ treated brain tissue. A strong proteome correlation was observed for control versus Aβ treated tissues. As a reference, two protein markers with strong correlation between control and Aβ treated tissies are highlighted (DLG4 (PSD95) and SYUA protein). A clear correlation deviation was observed for the two synaptic markers SMS and NEUG.

Next, we carried out protein extractions from Tg2576 mouse (age: 9months) brain tissue as well as human post-mortem AD and HC brains, using either TBS, formic acid (FA) or SDS extraction buffers, followed by immuno-precipitation with JD1 and downstream targeted MS analysis. IP-MS analysis of Tg2576 mice and human AD brains provided unbiased confirmation of the presence of JD1 positive Aβ assemblies in these brain samples (Supplementary Data Fig. 5a) and detection of these Aβ species was found to be increased using FA and SDS extraction buffers, when compared to the more soluble TBS fractions, which is an observation also reported by others (10). However, we cannot rule out that the use of harsh extraction buffers, such as FA, could eventually result in substantial dissociation of small oligomeric Aβ assemblies, therefore contributing to a decrease or even loss of the conformational epitope associated with JD1 binding.

**Figure 5.**
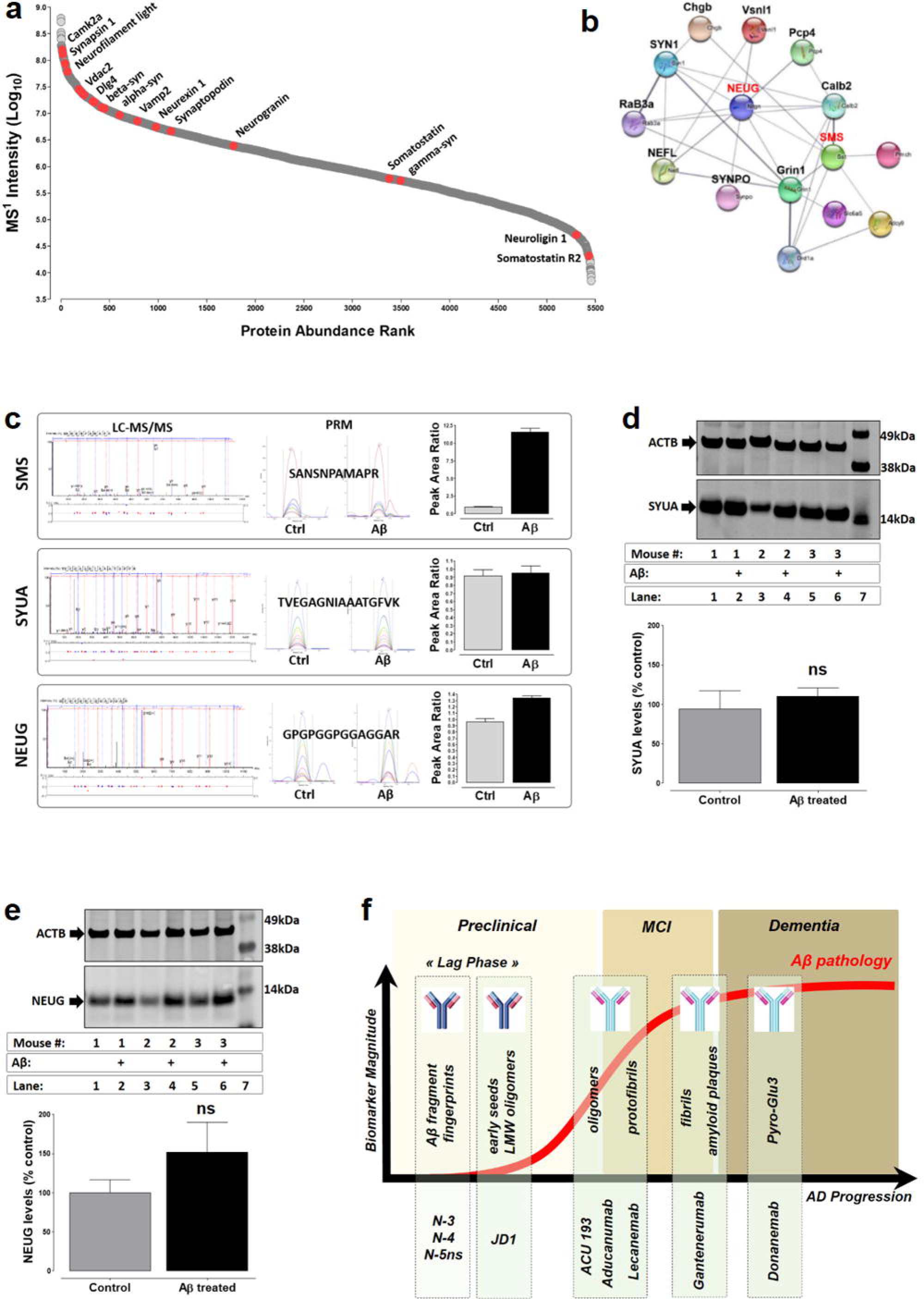
Acute Aβ neurotoxicity induces rapid changes in somatostatin and neurogranin levels in the mouse hippocampus. **a** Protein abundance analysis showing the relative abundance of different pre- and post-synaptic markers highlighted in red. **b** Selected zoom of the protein association network analysis (STRING), centred on neurograning (Neug) protein and showing an interaction with somatostatin (SMS) protein as well as a network with other proteins. (see Supplementary Data Fig. 8 for larger interaction network). **c** Summary of label-free, quantitative MS of the pre- and post-synaptic markers: somatostatin, alpha- synuclein and neurogranin (from top to bottom). Left; MS^2^ spectra of one proteotypic peptide monitored for SMS, SYUA and NEUG protein. Centre; MS^2^ fragment areas of selected MS^2^ transitions. Right; calculated relative protein areas derived from quantitative MS analysis. **d** Orthogonal validation of SYUA levels found in Aβ treated (lanes: 2, 4 & 6) and control (lanes: 1, 3 & 5) brain tissue. (lane 7: MWM) using WB. Below; Summary of actin normalized (ACTB) densitometry analysis of NEUG levels detected in Aβ treated (110.3±10, mean ± SD, n=3) versus control (94.3±23) (p=0.489, t-test, two-tailed) animals. (ns: not significant). **e** Orthogonal validation of NEUG levels found in Aβ treated (lanes: 2, 4 & 6) and control (lanes: 1, 3 & 5) brain tissue. (lane 7: MWM) using WB. Below; Summary of actin normalized (ACTB) densitometry analysis of NEUG levels detected in Aβ treated (152±38, mean ± SD, n=3) versus control (100±16.5) (p=0.082, t-test, two-tailed) animals. (ns: not significant). **f** Summary of neo-epitope (N3, N4 & N5ns) and JD1 antibody binding features, as well as earlier reported binding properties of different clinical antibodies. The diagram depicts a perspective of the current understanding of the pathological accumulation of Aβ assemblies (morphologies) and antibody molecules against these different Aβ entities: oligomers (ACU 193, aducanumab) and protofibrils (lecanemab), fibrils or amyloid plaques (gantenerumab, aducanumab) or modifications (pyro-E3) found within the N-terminal region of Aβ (donanemab). The combination of both, neo-epitope and JD1 antibodies may allow early interception of small oligomeric species prior to the accumulation of larger Aβ assemblies.

To further corroborate the presence of JD1 positive species in post-mortem human brain, we next, applied a sandwich ELISA (Supplementary Data Fig. 5b), using JD1 as capture antibody and a biotinylated form of 6E10 for detection of oligomeric Aβ species in HC (n=3) and AD (n=7) cortical brain tissue extracts. Our ELISA measurement revealed that, JD1 positive assemblies can be detected in all of the AD subjects, though, with considerable inter-subject variability (30-140pg / mg cortical brain extract), which may reflect the general variability in Aβ pathology found in AD subjects (Fig. 3d). Indeed, a large inter-subject variability of LMW Aβ assemblies was also observed by IP-WB (Fig. 3c). No JD1 positive species were detected in brain extracts from control subjects, however, the low number of subjects used here would not rule out the presence of low amounts of oligomeric Aβ in other, age matched control subjects or patients suffering from MCI (35).

Collectively, our brain imaging and screening studies strongly indicate that JD1 has preferential binding properties against small Aβ assemblies and that antibody binding may be gradually lost during the time course of on-pathway Aβ peptide aggregation. This may also explain the lack in JD1 binding to dense core plaques as well as the decrease in JD1 binding affinity following extended (≥2week) Aβ peptide aggregation (Supplementary Data Fig. 3b). Nevertheless, our findings do not rule out binding of JD1to protofibrillar or fibrillar structures, given that the Aβ binding epitope is sufficiently conserved and more importantly exposed, in these larger Aβ assemblies.

### LMW oligomers formed by fragments Aβ1-23 to Aβ1-25 significantly impair synaptic plasticity

Earlier reports showed that unlike full-length Aβ42, the truncated form of Aβ1-16 did not have any inhibitory effects on synaptic plasticity (36), whereas other reports indicate that the larger Aβ1-25 isoform, serves a sphingolipid-binding domain motif and is rapidly internalized by neuronal cells (37). This rapid peptide internalization may account for the acute neurotoxicity observed for some Aβ fragments. Therefore, to address the acute neurotoxic properties of truncated Aβ species, we decided to generate LMW oligomers using a mixture of the typically observed Aβ fragment fingerprints during aggregation. For this purpose, a mixture of the synthetic Aβ fragments Aβ1-23, Aβ1-24-NH2, Aβ1-25 and Aβ1-25- NH2 (50ng/ul) was incubated over-night followed by size exclusion chromatography (SEC) purification of LMW oligomeric assemblies that are recognized by JD1.

We identified JD1 positive Aβ species in the elution volume (13ml) corresponding to the mass of a theoretical molecular standard of ≥13.7kDa (Fig. 4a). WB analysis of the SEC fractions revealed a population of different Aβ assemblies of molecular weights ranging from

≥5kDa (dimers) to ≥ 6kDa (trimers), though we cannot rule out that these Aβ entities are the result of the SDS-PAGE analysis induced dissociation of larger Aβ assemblies. MS analysis of the JD1 positive fraction confirmed the identification of a mixture of all four Aβ isoforms present in the JD1 positive fraction (data not shown).

To set out a strategic approach in assessing the acute neurotoxic properties of LMW oligomers, we decided to incubate mouse hippocampal brain slices with the above described, SEC purified JD1 positive assemblies, followed by *in vitro* electrophysiological recordings of long-term potentiation (LTP) in the Schaffer collateral commissural pathway of the CA1 region in mouse hippocampal slices. Brain slices were incubated with 500nM JD1 positive Aβ assemblies or an equal volume of PBS buffer (controls), followed by baseline recordings for 20min prior to high frequency stimulation (HFS) and LTP recording during 60min. LTP measurements were paired in a way, where control and Aβ treatment was carried out on hippocampal tissues derived from the same mouse brain.

Hippocampal tissues treated with these small oligomers resulted in a statistically significant reduction (123.8 ± 9.05, n=5) (p≤0.01) in LTP measured at 1hr post-HFS, when compared to matched controls (190.6 ±22.94, n=5,) (Fig. 4b), which concurs with earlier reports showing an Aβ mediated attenuation of LTP *in vivo* and *in vitro* (5,17,18). IP-MS (JD1) analysis of the aCSF bath-solution showed that JD1 positive oligomers remained stable during the time course of LTP recordings (data not provided).

### LMW oligomers induce acute changes in somatostatin and neurogranin levels in the mouse hippocampus

Following LTP recordings, brain slices from control and Aβ treated hippocampal tissues were subjected to protein extraction, followed by downstream deep proteomic analysis to assess any putative changes in synaptic markers following acute treatment with LMW Aβ assemblies. We employed a relative quantitative proteomics approach using isobaric labelling with tandem-mass-tags (TMT) followed by off-gel isoelectric focusing fractionation (IEF) (Fig. 4c), to study the relative change in brain protein abundance and post-translational modifications (PTM) as a result of acute Aβ treatment.

Deep proteomic analysis of mouse hippocampal tissues allowed for unbiased identification of ≤5440 protein groups (FDR ≤1%) of which 6% of this proteome is associated with cellular signalling mechanisms (Fig. 4d-e, Supplementary Data Fig. 6-8). Overall, extended bioinformatics analysis of the mouse brain proteome did not reveal major changes in protein abundances in control versus Aβ treated tissues (Fig. 4d-e and Supplementary Data Fig. 9). This observation is in agreement with the general knowledge that *de novo* protein synthesis is mainly associated with late-phase LTP, consisting of several hours or days (38). We nevertheless found a statistically significant fold-change (-Log p = 2.84) of the pre-synaptic protein somatostatin (SMS) following Aβ treatment (Fig. 4d-e and Supplementary Data Fig. 9a&b). Interestingly, rapid hippocampal SMS release has been associated with acute stress in animals (39) and somatostatin is known to play a key role in regulating Aβ metabolism through modulating proteolytic degradation by neprilysin (40).

Next, a functional protein partnership search was carried out using the STRING database for the integration of both, known and predicted protein-protein interaction networks (physical or functional) to identify potential somatostatin interaction networks (Fig. 5a-b and Supplementary Data Fig. 8a-b). Active somatostatin interaction was found for the protein neurogranin (Neug / Nrgn), a well-established marker of neurodegeneration (41, 42), whereas neurogranin itself appears to form a network hub with different pre- and post-synaptic markers (Fig. 5b). Bioinformatics analysis revealed a small fold change (Log2 fold change = 0.58) for NEUG in Aβ treated tissue, however, this marker failed to show statistical significance (-Log p = 1.34) (Fig. 4d-e). The small change in NEUG levels following Aβ treatment is an intriguing observation because neurogranin expression is known to be regulated by synaptic activity (43) and memory encoding requires a rapid (≤ 15min) *de novo* synthesis of neurogranin in the mouse hippocampus (44). Moreover, neurogranin is known to play a key role in the regulation of metaplasticity, by lowering the threshold for LTP induction (45) and the Aβ oligomer induced inhibitory effect on LTP can be reversed by increasing neurogranin levels (46).

To confirm the significance of the differentially regulated levels found for somatostatin and neurogranin, we decided to analyse the abundance of these two synaptic markers using a targeted, quantitative MS approach known as Parallel Reaction Monitoring (PRM), which allows unbiased identification of low abundant species in complex sample matrices. Quantitative MS analysis revealed that, levels of SMS were indeed significantly increased in Aβ treated mouse brains when compared to control brain slices. Similarly, NEUG levels where approximately ≥ 30% increased in Aβ treated brains, whereas levels of the predominantly found presynaptic marker α-synuclein (SYUA) did not differ between both groups (Fig. 5c and Supplementary Data Fig. 9b&c). Orthogonal validation by WB of the pre- and postsynaptic markers of SYUA (Fig. 5d) and NEUG respectively (Fig. 5e), as well as homer1 and post-synaptic density protein (PSD95 / Dlg4) (Supplementary Data Fig. 9c), further corroborate the findings by MS. However, immuno-blot detection of the small, presynaptic protein somatostatin proved to be challenging (Supplementary Data Fig. 9c) and therefore highlights the advantage of implementing targeted and quantitative MS assays, for an unbiased identification and quantification of scarce proteins (Supplementary Data Fig. 10).

### Aβ oligomer mediated changes in mouse hippocampal phospho-proteome

It has been well established that Aβ accelerates hyperphosphorylation of tau protein, which in turn aggravates neuronal tauopathy (47). We therefore sought to identify putative changes in site-specific protein phosphorylation, as a result of acute Aβ treatment, using a titanium- dioxide (TiO_2_) bead enabled phospho-peptide enrichment approach, followed by LC-MS analysis. Overall, phospho-enrichment analysis allowed the identification of ≥7800 phospho- sites (FDR ≤1%), in control and Aβ treated hippocampal slices. Our target proteins of interest were primarily focused on well-established markers of synaptic plasticity, as well as proteins that have previously been reported to be susceptible to phosphorylation as a result of Aβ toxicity (48). Phospho-proteomic analysis revealed differential levels of site-specific phosphorylation of the calcium/calmodulin dependent protein kinase II alpha (CAMKIIa) protein, post-synaptic density protein (PSD95 / Dlg4), synapsin 1 & 2 (SYN1 & SYN2), synaptophysin (SYPH) as well as tau (TAU) protein in control versus Aβ treated brains (Supplementary Data Table II). In spite of the here reported MS identification of some differentially modified tau species found in acute Aβ treated brains, we were not able to confirm earlier reported tau phosphorylation *in vitro* at residues pS202 or pS416 using structurally different Aβ assemblies (49).

Collectively, our deep proteomic and phospho-proteomic analysis may further shed light on the acute Aβ mediated effects on synaptic plasticity. The observed, differentially regulated synaptic protein somatostatin and, to some extent neurogranin, is intriguing, and clearly merits a more thorough investigation in the future but is beyond the scope of the current study.

## Discussion

In this study, we provide analytical insight into the composition of LMW Aβ oligomers. We show that small Aβ assemblies of putative dimeric, trimeric and tetrameric nature consist of a combination of full-length Aβ42 as well as a repertoire of C-terminally truncated Aβ species, of which the latter may also independently seed to form SDS-stable, HMW oligomeric assemblies of 50kDa-200kDa mass range. Our unbiased MS identification of identical Aβ fragment fingerprints, together with the JD1 positive assemblies detected in post-mortem AD brains, further highlights the pathological significance of Aβ fragments; Aβ1-23 to Aβ1-25. Generally, our findings allow to draw strong parallels with earlier reported observations of typical Aβ fragment fingerprints (12) as well as the detection of a repertoire of different N- and, C-terminally truncated Aβ species in human CSF samples (36,49,50). The independent report of specific Aβ fragments found in human post-mortem AD brains as well as Down-syndrome (DS) subjects (51, 52) further corroborates the significance of the findings reported here. Nevertheless, it remains to be confirmed, if the here reported key findings mirror earlier reports on the inhibition of LTP, using similar LMW oligomeric assemblies (22) or, soluble ∼7kDa Aβ species isolated from post-mortem AD brains (9,10,15).

We provide compelling analytical proof, that the application of our recently developed neo- epitope antibodies against different truncated Aβ isoforms allows for a highly selective pull-down (IP) of SDS-PAGE stable assemblies from human post-mortem AD brains. These isolated Aβ assemblies strongly resemble the LMW Aβ species (4-7kDa) typically found following IP of formic acid induced dissociation of ADDLs. Our findings are in strong agreement with earlier reports indicating, that LMW Aβ oligomers consist of a repertoire of truncated Aβ species and full-length Aβ42 or Aβ40 (16, 53). However, earlier analytical attempts in identifying specific Aβ isoforms associated with LWM Aβ assemblies and, more specifically, the ∼4kDa and ∼7kDa Aβ species, have failed (16).

The lack in scientific and clinical consensus on the therapeutic efficacy of some clinical antibodies as well as the ongoing, heated debate on viable, oligomeric Aβ targets (54) (55), emphasizes the strong need for identifying highly specific Aβ targets for the development of next generation disease modifying-therapies. Recent studies corroborate the notion, that therapeutically targetable Aβ seeds exist during the lag phase of protein aggregation, and that acute administration of aducanumab in young mice led to a significant reduction of Aβ pathology in the mouse brain (1). Therefore, a successful clinical translation of specific Aβ target hotspots will allow a therapeutic interception of early Aβ seeds during the long, insidious lag phase of AD and prior to the accumulation of larger, metastable Aβ assemblies of protofibrillar or fibrillar nature (Fig. 5f). Improving antibody-binding properties for small Aβ assemblies may allow a selective targeting of early, pathological Aβ seeds during disease onset and therefore attenuate or even prevent the progression to severe dementia. The here reported monoclonal antibody binding features observed with JD1 represents a first example of a new class of future antibody molecules.

Our understanding of JD1 binding properties give rise to important Aβ target “loopholes” associated with earlier developed anti-conformational antibodies. Yet, it remains to be determined, if other antibody molecules, such as the anti-oligomer ACU 193 (3) or anti-protofibril antibody lecanemab, currently undergoing clinical validation (56, 57), provide similar binding properties for the here described LMW Aβ assemblies. Our observation on aducanumab’s decreased binding affinity for LMW oligomeric assemblies has not been reported before, and this finding is somehow surprising, because aducanumab’s binding epitope is known to span the N-terminal region of residues 3-6. Preferential aducanumab binding was reported for oligomeric and fibrillar Aβ species derived from full-length Aβ42 peptide (29), whereas oligomers derived from the N-terminally truncated forms of Aβ 5-42 to Aβ11-42 were no longer detected with aducanumab (58). It is conceivable, that the strong seeding propensity observed with fragments Aβ1-23, Aβ1-24 and Aβ1-25, as well as the C-terminally amidated forms of these fragments, involves rapid conformational changes associated with the amyloid driving segment of residues 16-25 and that JD1 binding is associated with a unique conformational structure found within this Aβ segment. Discrepancies in aducanumab binding properties have been previously observed, following immunization of a transgenic mouse model of AD, resulting in a clear reduction in amyloid plaques but a failure in clearance of vascular amyloid (CAA) (1, 29). Interestingly, it was reported that the deposition of specific Aβ isoforms appears to differ in subjects with vascular dementia, when compared with AD patients (51), which is in line with the positive immuno- staining found for fragment Aβ1-25 in CAA subjects (12).

Significant IHC staining of Aβ oligomers was reported using different conformation specific antibodies (3) with binding properties against large (≥80kDa) Aβ assemblies (59). The lack in JD1 staining of amyloid plaques may not appear surprising, given JD1’s binding specificity for LMW Aβ assemblies and the overall scarcity of Aβ oligomers reported in the human brain, which is an observation also made by others using different anti-oligomer antibodies (3,28,61). Nevertheless, our high-resolution array tomography clearly shows co-localization of JD1 positive species at post-synaptic terminals, which is in line with earlier reports, using different conformational specific antibodies (34).

Taken together, the selective binding profile found with JD1 could be a beneficial antibody feature, by decreasing the risk of dose-limiting side effects associated with passive immunization, as well as increasing the antibody’s bioavailability for rather early, neurotoxic oligomeric assemblies (61). Overall, the here reported binding properties of JD1 highlights the importance in further improving and developing novel antibody tools, with high target selectivity for small, pathological Aβ seeds or oligomeric entities derived thereof. To date, the development of highly specific anti-oligomeric antibodies remains a strong unmet medical need for future disease-modifying therapeutics in AD. Improvement of immuno-based target selectivity for acute, pathological Aβ species will inevitably provide early insight into the disease pathology. This in turn will serve as an indispensable diagnostic tool for evaluation of disease onset and progression, prior to the accumulation of clinical targets associated with pyro-Glu_3_ modification (Donanemab) (61) or large Aβ assemblies of protofibrilar (lecanemab) (57) and fibrillar structure (gantenerumab) (56) (Fig. 5f).

Monitoring changes in CSF and plasma Aβ42/40 ratios, as well as detection of early Aβ seeds, together with other, clinically well-established biomarkers of neurodegeneration (e.g. p-Tau, Nfl, Ngrn), may overall improve the diagnostic accuracy for individuals suffering from mild cognitive impairment, who may benefit from early therapeutic intervention with future disease modifying therapies. This could help to attenuate the pathological progression to severe dementia and overall improve the quality of life in patients in the long run.

## Author contributions

AWS conceived the project, designed and directed the study. CEH and BG planned and carried out LTP studies. AWS performed mass spectrometry and biochemical analysis. JR and TS planned and carried out IF and array tomography studies on human brain tissue. GM and JSD carried out IF studies on Tg mouse brain. DD performed electron microscopy. VS carried out *C. elegans* experiments and analysis. MPF performed histological examination of human brain tissue. NR and AWS carried out ELISA measurements. BTH and MPF provided biological samples. AWS and CEH wrote the paper and all authors read and commented on the manuscript.

## Acknowledgments

This work was funded by the “Enable Grant”, kindly awarded by Dr. André Catana from the Technology Transfer Office (TTO) as well as the Catalyze4Life Innovation grant by Prof. Bart Deplancke and Dr. Kostas Kaloulis of the Ecole Polytechnique Fédérale de Lausanne (EPFL). We thank Dr. Adam Swetloff (TTO) for the constructive and strategic discussions on the project development as well as the support by the whole Life Sciences Faculty. The project was partially funded using internal funding of the EPFL. Human brain sample preparation and ELISA development was supported by the ADRC grant P30AG062421.We are grateful to the Roland Bailly Foundation (Geneva, Switzerland) for the funding of the MS instrument. We extend our gratitude to Dr. David Hacker (Protein Production and Structure Core Facility (EPFL) for his help with antibody production. Prof. Carmen Sandi, Laboratory of Behavioural Genetics (EPFL) for kindly providing us with antibodies for the study as well as Dr. Jose Sanchez-Mut, Laboratory of Neuroepigenetics (EPFL, Prof. J. Gräff lab), providing us with the reversed sequence of Aβ42-1 peptide and Prof. Hilal Lashuel (EPFL) for the Aβ40-arc mutant peptide. We thank Dr. Pamela Valdés (LEN, EPFL) for her support with IF imaging. TSJ and JR are funded by the UK Dementia Research Institute which receives its funding from DRI Ltd, funded by the UK Medical Research Council, Alzheimer’s Society, and Alzheimer’s Research UK, and the confocal used for IHC/AT was funded by Alzheimer’s Research UK (ARUK-EG2016A-6) and a Wellcome Trust Institute Strategic Support Gant.

## Materials and Methods

### Aβ peptide preparation

Full-length wild type (wt) Aβ peptides Aβ1-40 and Aβ1-42 Aβ (BACHEM, Switzerland) were dissolved in 1,1,1,3,3,3-Hexafluoro-2-propanol (HFIP) at a concentration of 1mg/ml, followed by a 10-min sonication to break any preformed aggregates. HFIP solution was evaporated under a ventilated fume hood by applying a light stream of N_2_ gas. The HFIP film containing the Aβ peptide was either directly re-suspended in 100% DMSO and further diluted to 1% DMSO in a new buffer or stored dry at −80 °C until use. Aβ peptide fragments comprising of residues: 1-23, 1-24-NH_2_, 1-25, 1-25-NH_2_ (purity of ≥ 97%) were purchased from GenicBio Ltd. (Shanghai, China) and Aβ1-42 and Aβ1-40 were purchased from Bachem (Schwitzerland), Aβ40-Arc was kindly provided by Prof. Hilal Lashuel (EPFL) and reverse peptide Aβ42-1 was kindly provided by Prof. Gräff (EPFL). Recombinant alpha-synuclein protein was a gift from rPeptide (GA, USA).

### In vitro Aβ peptide aggregation studies

Aggregation kinetics were carried out as reported by (Ladiwala et al.). Briefly, HFIP dried Aβ peptide films were solubilized in 50mM NaOH, sonicated and further diluted to 5mM with PBS (or 30mM Tris, 150mM NaCl) to a final Aβ concentration of 0.05-0.1 mg/ml for truncated Aβ fragments and full-length peptides of Aβ1-40, 0.1 mg/ml mutant Aβ1-40 Arc, and 0.05 mg/ml Aβ1-42. Aβ preparations were then centrifuged (22,000 x g) for 30min at 4°C and the supernatant (90-95% volume) was recovered and transferred into a new Eppendorf tube. Supernatant samples were then incubated at 37°C and left for spontaneous aggregation during either 0-20 hrs (short-term) or 1-5 days (intermediate). Aggregated sample aliquots were drawn at different time points and either analyzed immediately using MS and/or DB or snap frozen in liquid nitrogen and stored at -80°C. Typically, one microliter was spotted onto a nitrocellulose membrane for dotblotting (corresponding to either 25, 50ng to100ng / spot) or mixed with alpha-cyano matrix for MALDI-TOF/TOF analysis.

### Digestion of Aβ peptides

Proteolytic digestion using LysN (2ng/ul) was performed overnight at 37 °C in 50 mM ammonium bicarbonate, pH 10 (LysN buffer). For in-gel digestions, peptides were extracted from gels and concentrated by Speed-Vacuum prior to LC-MS/MS measurements. Immunoprecipitated samples (IP), were reduced and alkylated followed by in-solution digestion at 37°C using standard LysN buffer (50ul) and approximately 70ng of LysN. Dried samples were resuspended in 5% DMSO and 2.5% FA followed by LC-MS/MS, LC-MRM and PRM analysis.

### Matrix assisted Laser Desorption/Ionization Time-of-flight Mass Spectrometry (MALDI TOF/TOF)

Aliquots (2 μl) of samples were used for MALDI-TOF/TOF MS (ABI 4800 model, Applied Biosystems) measurements. Matrix solution of α-cyano-4-hydroxycinnamic acid (7 mg/ml in ACN/0.1% TFA (1:1, v/v)) was used for sample deposition. The sample (1 μl) was mixed with 1 μl of matrix solution and then 1 μl of this mixture was deposited in duplicates on the target plate and allowed to air dry. Samples were analyzed in reflectron positive mode.

### Sandwich ELISA for detection of Aβ oligomers

JD1 antibody was adsorbed to high protein binding Nunc-Immuno MaxiSorp 96-well plates (Nunc, Roskilde,) at 4°C for 4hrs, followed by a blocking step for 1hr using a 5% BSA in PBST (Tween 20: 0.1%) solution. Synthetic, oligomeric preparations of Aβ1-24 or Aβ1-25 standards or, human TBS and FA brain extracts were incubated at 4°C over-night to allow for binding to the capture antibody JD1. Detection of Aβ species was achieved using the mouse monoclonal HRP-conjugated 6E10 (Biolegend) antibody at a dilution of 1:1000. Signals were measured at an absorbance at 450nm using a Tecan Infinite F500 reader. All samples were analyzed in triplicates.

### Mouse hippocampal tissue extraction, TMT labelling and LC-MS analysis

Mouse hippocampal proteins from PBS control and Aβ oligomer (JD1 positive) treated brain slices were extracted with 2% SDS (in PBS) containing protease and phosphatase inhibitors (Roche, Switzerland) followed by centrifugation at 14000 x g for 15min to precipitate any insoluble material. Prior to proteolysis, samples were cleared of excess SDS using detergent removal spin columns (Cat# 87777, Pierce, Switzerland) followed by total protein content normalization using a standard Bradford protein assay. Samples were reduced and alkylated followed by overnight digestion at 37°C using LysC / Trypsin proteases (Promega) and isobaric labelling using tandem mass tags (TMT) as described elsewhere. Briefly, the TMT reagent (0.8 mg, Thermo Fisher Scientific) was dissolved in acetonitrile, and 10 µL of the reagent was added to 30 µL of peptides along with anhydrous ACN to give a final concentration of 30% v/v. The peptides were then pooled after incubating at room temperature for 1h followed by C18 stage tip clearing and drying in a speed vacuum.

For Off-gel fractionation, samples were suspended in a 4M urea buffer containing ampholytes and subjected to off-gel IEF fractionation using a 24 well strip (pH 3-10) (Agilent, 3100 Off- gel fractionator). Following IEF, fractions were recovered and samples were cleared with C18 stage tips followed by speed vacuum drying and storage at -20°C until LC-MS analysis.

### LC-MS/MS analysis

Mass spectrometry (MS) analysis was performed on an Orbitrap Exploris 480 mass spectrometer (Thermo Fisher Scientific) coupled to a nano-UPLC Dionex pump. For liquid chromatography (LC) MS/MS analysis, trypsin / LysC digested samples were resuspended in 30-60ul of a mobile phase A solution (2% ACN / water, 0.1% formic acic) and then separated by reversed-phase chromatography using a Dionex Ultimate 3000 RSLC nanoUPLC system on a home-made 75 µm ID × 50 cm C18 capillary column (Reprosil-Pur AQ 120 Å, 1.9 µm) in-line connected with the MS instrument. Peptides were separated by applying a non-linear 150min gradient ranging from 99% solvent A (2% ACN and 0.1% FA) to 90% solvent B (90% ACN and 0.1%FA) at a flow rate of 250nl/min. Full-scan MS spectra (300−1500 m/z) were acquired at a resolution of 120’000 at 200 m/z. Data-dependent MS/MS spectra were recorded followed by HCD (higher-energy collision dissociation) fragmentation on the ten most intense signals per cycle (2 s), using an isolation window of 1.4 m/z. HCD spectra were acquired at a resolution of 60’000 using a normalized collision energy of 32 and a maximum injection time of 100 ms. The automatic gain control (AGC) was set to 100’000 ions. Charge state screening was enabled such that unassigned and charge states higher than six were rejected. Precursors intensity threshold was set at 5’000. Precursor masses previously selected for MS/MS measurement were excluded from further selection for a duration of 20 s, and the mass exclusion window was set at 10 ppm. Theoretical isotopic envelopes of Aβ peptides were drawn using the isotope calculator tool in Data Explorer Software (version 4.9). The mass spectrometry data generated from this study have been deposited to the ProteomeXchange Consortium (http://www.proteomexchange.org) via PRIDE partner repository with the dataset identifier: PXD033934 (10.6019/PXD033934).

### Targeted and quantitative MS analyses

Parallel reaction montitoring (PRM) analyses were performed as reported earlier (Sathe et al) using an Orbitrap Exploris 480 mass spectrometer (Thermo Fisher Scientific) coupled to a nano-UPLC Dionex pump. Peptides from selected target proteins, SST, NEUG, SYUA were monitored using a targeted inclusion list. The parameters used for the PRM analysis were set as follows: Orbitrap resolution of 35,000, AGC target value of 5 x 10 5, injection time of 200 ms. A precursor target isolation window of 2 m/z was applied and a normalized collision energy of 35 was employed for MS/MS fragmentation. The PRM method was carried out in an unscheduled mode and samples were analysed in duplicates. PRM data were analyzed using the Skyline software package (MacCoss Lab). Acquired raw files were uploaded to Skyline and peak integration was verified manually.

### MS raw data processing and database searches

PEAKS Studio X+ Pro (Bioinformatics Solutions Inc.) software was used for data processing. The raw MS data files were imported into PEAKS Studio software using the following parameters for the database search: For protein identification the UniProt / Swiss-Prot mus musculus database (ID: UP000000589) combined with a decoy database was used. For peptide identification the following settings were used: Enzyme: Trypsin, missed cleavages: 2 precursor mass tolerance: 10 ppm, fragment mass tolerance: 0.2 Da, minimum charge: 2, maximum charge: 5, fixed modifications: Carbamidomethyl (C), variable modifications: Oxidation (M), phosphorylation (STY). False discovery rate (FDR) was calculated based on the target/decoy database and peptides as well as proteins with FDR threshold of ≤ 1% were chosen as true positive hits.

MaxQuant searches: Label-free and TMT-labelled mouse brain proteomes were processed by MaxQuant as reported earlier (Tyanova 2016). Briefly, MS raw files were processed using the peptide identification algorithm of the Maxquant software (ver. 1.5.3.12) using the Andromega search engine. TMT correction factors were updated according to the values provided manufacture. MS/MS searches were performed with the following parameters: Peptides had to have a minimum length of six amino acids to be considered for identification. Carbamidomethylation was set as a fixed modification, while oxidation of methionine and protein N-terminal acetylation were set as variable modifications. The enzyme specificity was set to trypsin and allowed up to two miscleavages. The UniProt / Swiss-Prot mus musculus database (ID: UP000000589) combined with a decoy database was used. A false discovery rate (FDR) of ≤1% was applied at the proteins and peptide level. Quantitative data analysis was performed using Skyline (version 21.1.0.278, MacCoss lab, Uni. Washington, USA), an open source software tool application for quantitative data processing and proteomic analysis. All integrated peaks were manually inspected to ensure correct peak detection and integration.

Bioinformatic analysis was carried out using the Perseus (v.1.6.15.0) software environment and compared with the bioinformatics output generated with the PEAKS Studio X+ Pro software to identify any putative discrepancies due to the analytical stringencies or search engine algorithms used in both workflows. Volcano plots were generated using the -log10 of the p-value on the y-axis versus the mean fold-difference of the log2 transformed quantitative data on the x-axis. A paired t-test with ‘Benjamini-Hochberg FDR’ method was applied for multiple hypothesis corrections. Significance curves in the volcano plot corresponded to an S0 value of 0.5 and an FDR cut-off of 0.05. Protein interaction networks were analyzed and plotted using the open source platforms: STRING DB (http://string-db.org) (Jensen et al.) and Cytoscape (v.3.9.0) software (Shannon et al).

### SDS-PAGE and Immunoblotting

Where indicated in the results, samples were mixed using either standard SDS Lämmli sample buffer and heated at 80°C for 5 min prior to loading onto gels (Novex Nupage 4-12% Bis-Tris gels, 1mm, 15 well, Invitrogen) and ran with Novex Nupage MES-SDS buffer. PAGE separated samples were electroblotted using a constant voltage of 30V (90min) onto nitrocellulose (0.22µm) membranes. Following transfer, membranes were boiled in PBS for 2min in a microwave oven (800W) and then blocked for 1h at room temperature under constant rocking using Odyssey blocking buffer (Li-COR Biosciences, Bad Homburg, Germany) diluted 1:3 in PBS. Following blocking, membranes were incubated at 4 °C with constant rocking overnight using the commercially available mouse monoclonal antibodies 6E10 and 4G8 (0.5µg /ml) (Enzo, Life Sciences, Switzerland). Membranes were washed four times with PBS-Tween (PBS containing 0.1% Tween 20), followed by incubation with a goat anti-rabbit or anti-mouse secondary IgG antibody (highly cross-adsorbed) (dilution, 1:5000) conjugated to Alexa Fluor 680 or 800 and scanned in a Li-COR scanner at a wavelength of 700 nm and 800 nm respectively. Densitometric analysis of WB band intensities was carried out using image J software v.1.43 (NIH).

### Dot blotting

Typically, 1ul samples were spotted onto a nitrocellulose membrane, which corresponded to a total peptide load of 25ng (Aβ1-40) and (AB1-42) unless otherwise stated in the Fig.ures. Samples were left to dry for 15min followed by blocking of the membrane (30min) with 5% BSA in PBS solution. Membrane strips were incubated with primary antibodies overnight at 4°C on a shaker. For JD1 probing, membranes were incubated 2-4hrs at RT or over-night at 4°C followed by revelation with a goat anti-mouse, secondary Alexa-flour conjugated antibody. Neo-epitope antibodies N3, N4, N5s N5ns and N5NH2 were applied as described earlier (12). Identical solutions and revelation procedures with secondary antibodies were used as described for immunoblotting above.

### Immunoprecipitation of Tg mouse and human brain tissue

Tg mouse (9-12 months) (Tg2576) and cortical human post-mortem brain tissue samples were extracted using either TBS, 1% SDS (in TBS) or formic acid (70-90%FA) as reported earlier Rudinskiy et al. (12) Briefly, tissue samples were homogenized (20 strokes on ice) in TBS with protease inhibitors (Roche, Switzerland) using a Teflon homogenizer. Samples were then subjected to centrifugation (150,000g) during 45min and supernatant was recovered as the TBS soluble fraction. Protein pellets were subjected to an additional extraction using either SDS (1%) or FA (70%) followed by centrifugation. Typically, pellets were extracted with 70-90% FA overnight at 4°C (to minimize formylation adducts) followed by centrifugation. SDS fractions were diluted to ≤0.1% SDS final concentration and FA fractions were neutralized to pH 7.5 with 5M sodium hydroxide (NaOH) solution prior to IP. All samples were initially depleted of endogenous IgG’s using a mixture of protein A&G agarose beads (Roche AG, Switzerland). Typically, 3-5ug/ml rabbit polyclonal antibody (N3, N4, N5ns & N5NH2), 5ug/ml of JD1 (IgG) or 3-5ug/ml of 6E10 or 4G8 mouse monoclonal was used for IP overnight under continues rotation (4rpm/min) at 4°C. Samples were eluted with 40% ACN /H_2_O & 0.1% TFA for downstream MS analysis and dried in a speed vacuum or, boiled in Lämmli buffer for WB analysis. IP’ed samples were either directly analysed by WB or MALDI-TOF/TOF or further digested overnight using LysN proteolysis prior to LC-MS/MS analysis as outlined above.

### Animals

Hippocampal slices were prepared from 8 to 10 week old c57/black 6 mice (c57BL6), obtained from Charles River UK. Mice were housed in the biomedical facility, University College Dublin with a 12 h light/dark cycle with food and water ad libitum. All experiments were conducted under license from the Dept. of Health, Ireland, Directive 63/2010/EU.

### Hippocampal Slices

Parasagittal hippocampal slices, were prepared using a vibrotome (Leica VT1000S) as described previously (62, 63) using ice cold cutting solution (mM) NaCl 87; NaHCO_3_ 25; Glucose 25; Sucrose 75; KCl 2.5; NaH_2_PO_4_ 1.25; CaCl_2_ 0.5; MgSO_4_ 7; bubbled with 95% O_2_/5% CO_2_ (carbogen). Slices (400-µm thick) were transferred to a holding chamber containing aCSF (mM) NaCl 119; NaHCO_3_ 26.2; Glucose 11 mM; KCl 2.5; NaH_2_PO_4_ 1; CaCl _2_ 2.5; MgSO_4_ 1(also used for recording); bubbled with carbogen and were allowed to recover at room temperature for at least 90 min Slices were transferred to a recording chamber, secured by means of a harp with fine nylon strings and perfused with recording aCSF at a rate of 5 ml/min maintained at 28–30 °C for the duration of all experiments. A horizontal puller (DMZ universal puller, Germany) was used to makerecording electrodes from borosilicate capillary glass (GC150 F-10, Havard apparatus), Electrodes (2–5 MΩ) were filled with recording aCSF. The Shaffer-collateral pathway was stimulated using a mono- polar electrode (FHC, Bowdoin, USA) at 0.033 Hz (duration: 100 µs), the return electrode was a silver/silver chloride wire placed in the recording bath. Extracellular field excitatory post synaptic potentials (fEPSPs) were recorded in the CA_1_ *stratum radiatum* and paired stimuli were delivered with an inter-stimulus interval of 50 ms in order to monitor paired pulse facilitation (PPF). The voltage signal was filtered at 5 kHz and stored for off-line analysis using a personal computer interfaced with a Digidata 1440 and pclamp 10. Signals were amplified by a HS2A headstage (Molecular Devices, USA) connected to an Axoclamp 2B system (Molecular Devices, USA) and a Brownlee 410 Precision preamplifier. The stimulus voltage was adjusted to evoke a fEPSP that was 40–50% of the maximal response (maximum fEPSP just prior to the formation of a spike caused by cell firing) for the duration of all experiments. When a stable baseline had been recorded for 20 min a Master 8 (AMPI) timer was used to deliver two trains of high frequency stimuli (HFS), 100 Hz for 1 s, with an inter-train interval of 30 s. Following the application of HFS, the synaptic response was recorded for a further period of 60 min. All results are presented as mean ± SEM. The “*n*” numbers quoted refer to the number of slices used. Control and test experiments in a given section were conducted on the same day on slices from the same animal.

### Development of monoclonal antibody against LMW Aβ oligomers

Briefly, mice were immunized with the linear sequence of Aβ1-24 (CONH2) as well as an aggregated form of Aβ1-24 peptide. Eight weeks following immunization, sera were tested by ELISA for antibodies against fresh (linear) and oligomeric Aβ species. Mice testing positive for anti-Aβ antibodies were then given booster injections of 50 ug of an oligomeric form of Aβ1-25. IgM type antibodies consisted of the predominantly identified type of anti- Aβ antibodies found following this protocol. Antibody type switching and grafting from IgM to IgG was then carried using recombinant technologies. All work was carried out by Eurogentec SA, (Liege, Belgium).

### Immunofluorescence and array tomography imaging of human post-mortem brain tissue

Samples from the inferior temporal gyrus of 4 AD patients was prepared for array tomography as described previously (12, 34). Brain samples were obtained with ethical approval via the University of Edinburgh Sudden Death Brain Bank and the Alzheimer Scotland Brain and Tissue Bank. This study was reviewed and approved by the Edinburgh Brain Bank ethics committee and the Academic and Clinical Central Office for Research and Development, a joint office of the University of Edinburgh and NHS Lothian (approval number 15-HV-016). The Edinburgh Brain Bank is a Medical Research Council funded facility with research ethics committee (REC) approval (16/ES/0084). Briefly, tissue was collected at autopsy, fixed for 3 hours in 4% paraformaldehyde with 2.5% sucrose, dehydrated through a series of ethanols, and embedded in LR White resin. Ribbons of 70nm serial sections were cut with an ultracut microtome (Leica). Ribbons were stained with, primary antibodies OC (Sigma AB2286, 1:200), JD1 (IgG, 1:200), and PSD95 (Synaptic Systems 124 014; 1:50) and detected with anti-mouse IgG alexa fluor 594, and anti-rabbit alexa fluor 405, and anti-guinea pig alexa fluor 647 (all at 1:50). Images from 5 serial sections were acquired on a Leica TSC SP8 confocal with a 63x oil objective. Individual images were made into stacks, aligned, and segmented with custom software that can be found at https://github.com/Spires-Jones-Lab For immunofluorescence staining of 4 μm paraffin sections from formalin fixed tissue, sections were de-waxed, rehydrated, and stained with amyloid beta antibodies JD1 (1:500) and OC (1:1000) and secondaries donkey anti-rabbit 647 and donkey anti-mouse 594. Thioflavin S counterstain was used to label fibrils (0.01%). Image stacks were acquired on a Leica TSC SP8 confocal with a 63x objective.

### Transmission Electron Microscopy (TEM)

Aβ peptide (5μl) samples were deposited on Formvar-coated 200-mesh copper grids (Electron Microscopy Sciences, Hatfield, PA). Grids were washed with two drops of double-distilled H_2_O and stained with two drops of freshly prepared 1% (w/v) uranyl formate (Electron Microscopy Sciences). The samples were analysed at room temperature using a Tecnai Spirit (FEI, The Netherlands) electron microscope equipped with a LaB6 filament and operated at an acceleration voltage of 80 kV. Images were recorded with an Eagle (FEI, The Netherlands) camera 4098 X 4098 pixels. Representative images of the main Aβ structures were taken at different magnifications ranging from 30kX to 50kX.

### C. elegans strain

C. elegans overexpressing Aβ42 peptide were developed as reported before (24). Briefly, c. elegans were cultured at 20 °C on nematode growth medium (NGM) agar plates seeded with E. coli strain OP50 unless stated otherwise. The GMC101 strain was used in this study and was provided by the Caenorhabditis Genetics Center (University of Minnesota).

