## Supplemental Figures for "Elucidation of Amyloid-Beta’s Gambit in Oligomerization: Truncated Aβ fragments of residues Aβ1-23, Aβ1-24 and Aβ1-25 rapidly seed to form SDS-stable, LMW Aβ oligomers that impair synaptic plasticity"

[1]Conway Institute of Biomolecular and Biomedical Research, University College of Dublin (UCD), Ireland.
[2]UK Dementia Research Institute and Centre for Discovery Brain Sciences at The University of Edinburgh, Edinburgh, Scotland.
[3]Interdisciplinary Centre for Electron Microscopy (CIME), EPFL, Switzerland.
[4]Histology Core Facility, Ecole Polytechnique Fédérale de Lausanne (EPFL), Switzerland.
[5] Healthy Longevity Translational Research Programme, Yong Loo Lin School of Medicine & Department of Biochemistry, National University of Singapore.
[6]Harvard Medical School, Mass General Hospital, Massachusetts Boston, USA
[7]Massachusetts General Hospital and Harvard Medical School, Massachusetts General Institute for Neurodegenerative Disease, Charlestown, MA, USA.
[8]Proteomics Core Facility, Ecole Polytechnique Fédérale de Lausanne (EPFL), Lausanne, Switzerland.
[+] Contributed equally

*To whom correspondence should be addressed:

Adrien W. Schmid, PhD.
Life Sciences (SV), PTP-PCF, Station 15, EPFL
1015 Lausanne, Switzerland


| No | ID | Type | Age | Gender | Race | PMI | CLDX | Disease Duration | NPDX | B&B; CERAD | ELISA | DB | WB | IP | IHC |
|---|---|---|---|---|---|---|---|---|---|---|---|---|---|---|---|
| **1** | 795 | brain | 71 | F | W | 8 | AD | | AD | VI; C | x | | x | x | |
| **2** | 829 | brain | 88 | F | W | 20 | AD | 12 years | AD | VI; C | x | | x | x | |
| **3** | 835 | brain | 76 | F | W | 21 | AD | | AD, CAA | II; C | x | | x | x | |
| **4** | 1035 | brain | 80 | F | W | 48 | HC | | C | I | x | | x | x | |
| **6** | 728 | brain | 93 | M | W | 93 | AD | 25 years | AD | V;A | x | | x | x | |
| **7** | 748 | brain | 78 | F | W | 20 | AD | 17 years | AD | V; C | x | | x | x | |
| **8** | 1000 | brain | 76 | M | W | 48 | HC | | C | I | x | | x | x | |
| **9** | 1346 | brain | 80 | F | W | 54 | HC | | C | | x | | x | x | |
| **10** | 331 | brain | 90 | F | W | 22 | AD | 7 years | AD | C | x | | x | x | |
| **11** | NA | plasma | 54 | M | W | | HC | | | | | x | | | |
| **12** | SD014/15 | brain | 85 | F | unknown | 45 | AD | unknown | AD | VI | | | | | x |
| **13** | SD010/16 | brain | 85 | M | unknown | 91 | AD | unknown | AD | VI | | | | | x |
| **14** | SD039/17 | brain | 89 | F | unknown | 96 | vasc. dementia | unknown | AD | VI | | | | | x |
| **15** | SD028/19 | brain | 83 | F | unknown | 95 | dementia | unknown | AD | VI | | | | | x |
| **16** | SD034/17 | brain | 61 | F | unknown | 69 | dementia | unknown | AD | VI | | | | | x |

**Supplementary Table I:** Subject demographics of human brain tissue samples and plasma sample used in the study. Demographic information of the subjects used for the biochemical analysis (WB, IHC and IP-MS). Abbreviations: M, male; F, female; W, white; PMI, post-mortem intervals; CLDX; clinical diagnosis NDPX, neuropsychological diagnosis; B&B, Braak stage; CERAD, consortium to establish a registry for AD; AD, Alzheimer's Disease; HC, human control.

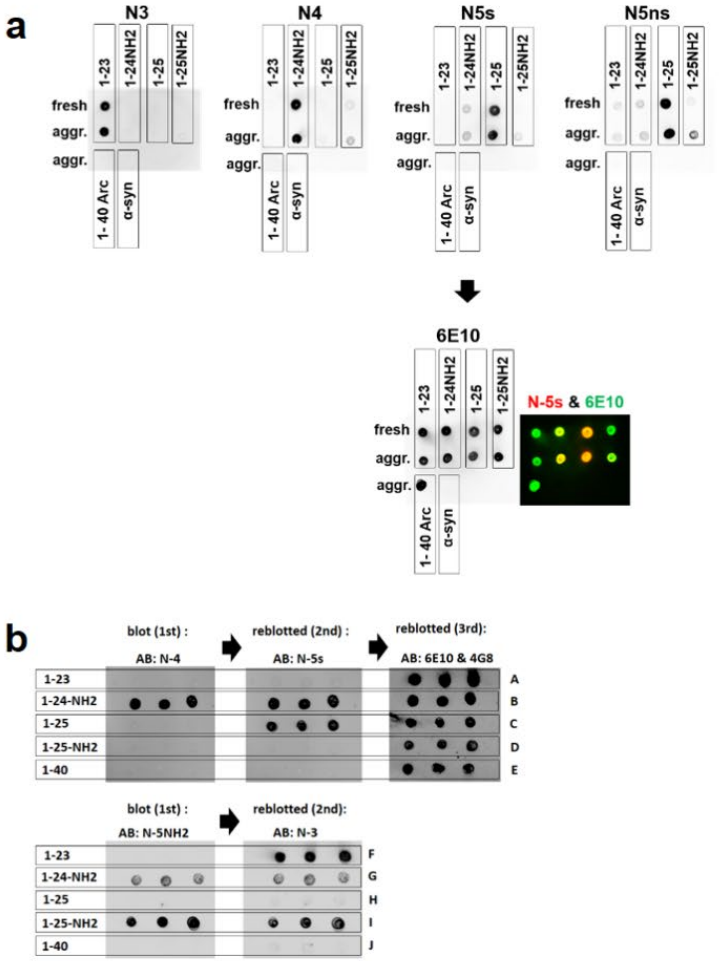


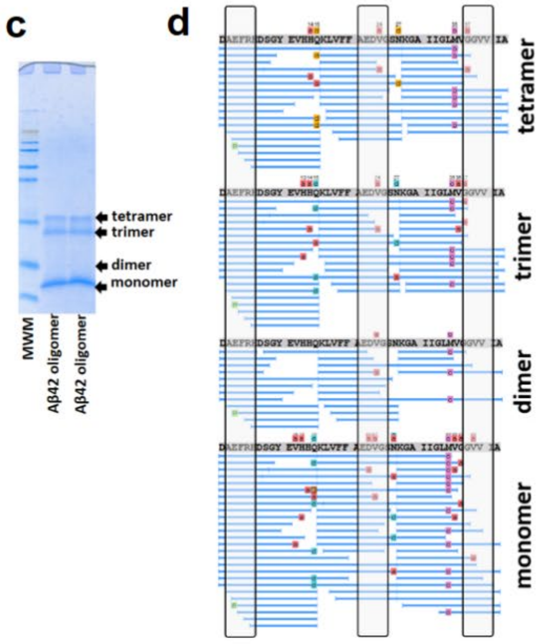


**e**

| Fragment | Monomer | Dimer | Trimer | Tetramer | Modification |
|---|---|---|---|---|---|
| 1-42 | x | x | x | x | |
| 1-40 | | | | | |
| 1-39 | | | | | |
| 1-38 | | | | | |
| 1-37 | | | | | Amidationn G37 |
| 1-36 | | | | | Amidation V37 |
| 1-28 | | | | | |
| 1-25 | x | x | x | x | |
| 1-25-$NN_2$ | | | | | amidation G25 |
| 1-24 | x | x | x | x | |
| 1-24-$NH_2$ | x | x | x | x | amidation V24 |
| 1-23 | x | | | | |
| 1-23-$NH_2$ | x | | | | Amidation D23 |
| 1-22 | | | | | |
| 1-20 | | | | | |
| 1-19 | | | | | |
| 1-18 | | | | | |
| 1-17 | | | | | |
| 1-15 | | | | | |
| 1-15 | x | | x | x | Deamidation Q15 |
| 1-14 | | | | | Amidation H14 |
| 1-13 | | | | | Amidation H13 |
| 1-12 | | | | | |
| 1-11 | | | | | |
| 12-24-$NH_2$ | | | | | amidation V24 |
| 12-25 | | | | | |
| 2-x | | | | | |
| 3-x | x | x | x | x | |
| 3-x | x | x | x | x | pyroE3 |
| 7-x | | | | | |
| 8-x | | x | x | | |
| 17-x | x | x | x | x | |
| 18-x | x | x | x | x | |

**Supplementary Data Figure 1: a** Neo-epitope antibody binding validation using dot blotting showing the binding selectivity of different antibodies against the truncated Aβ species in the soluble (freshly prepared) and aggregated peptide state. No antibody binding was observed against the aggregated form of the arctic mutant Aβ40 or alpha-synuclein. **b** Nitrocellulose membranes were prepared with rows of different Aβ peptide fragments (100ng/ spot, 3 replicates), rows A): Aβ1-23, B) Aβ1-24-NH2, C) Aβ1-25, D) Aβ1-25-NH2 and E) Aβ1-40. Membranes were blocked with 2% BSA during 1hr and then incubated for 2hrs with the rabbit polyclonal antibody N-4 (0.5ug/ml) (left) followed by a sequential (reblotting) of the membrane using a different neo- epitope antibody N-5s (center) and a final reblotting (3rd) with the commercial antibodies 6E10 and 4G8 (right). High specificity was observed for antibodies N-4 and N-5s showing binding to their specific antigens of C-terminal endings of V24-NH2 (amidated) and G25 respectively. A third and final reblotting of the membrane, using antibody 6E10 detection of all Aβ peptide fragments. ***Bottom panel;*** blots were prepared as above but using different antibodies for probing. left, membrane was probed using antibody N-5NH2 against the amidated form of C-terminal G25-NH2. N-5NH2 showed high specificity for G25-NH2 with some cross-reactivity (≥5%) for the shorter peptide fragment (1-24) with an amidated C-terminal at position V24-NH2. Antibody N-5NH2 does not bind the C-term carboxy form of fragment Aβ 1-25 (G25) (row: H). Reblotting the same membrane with antibody N-3 revealed high binding specificity against fragment Aβ 1-23 (row: F). **c - e** Summary of LC-MS analysis of gel extracted Aβ monomers to tetramers and the identification of Aβ sequences (PEAKS Studio) found in the different bands. A summary of major Aβ species identified in these bands are provided (e).

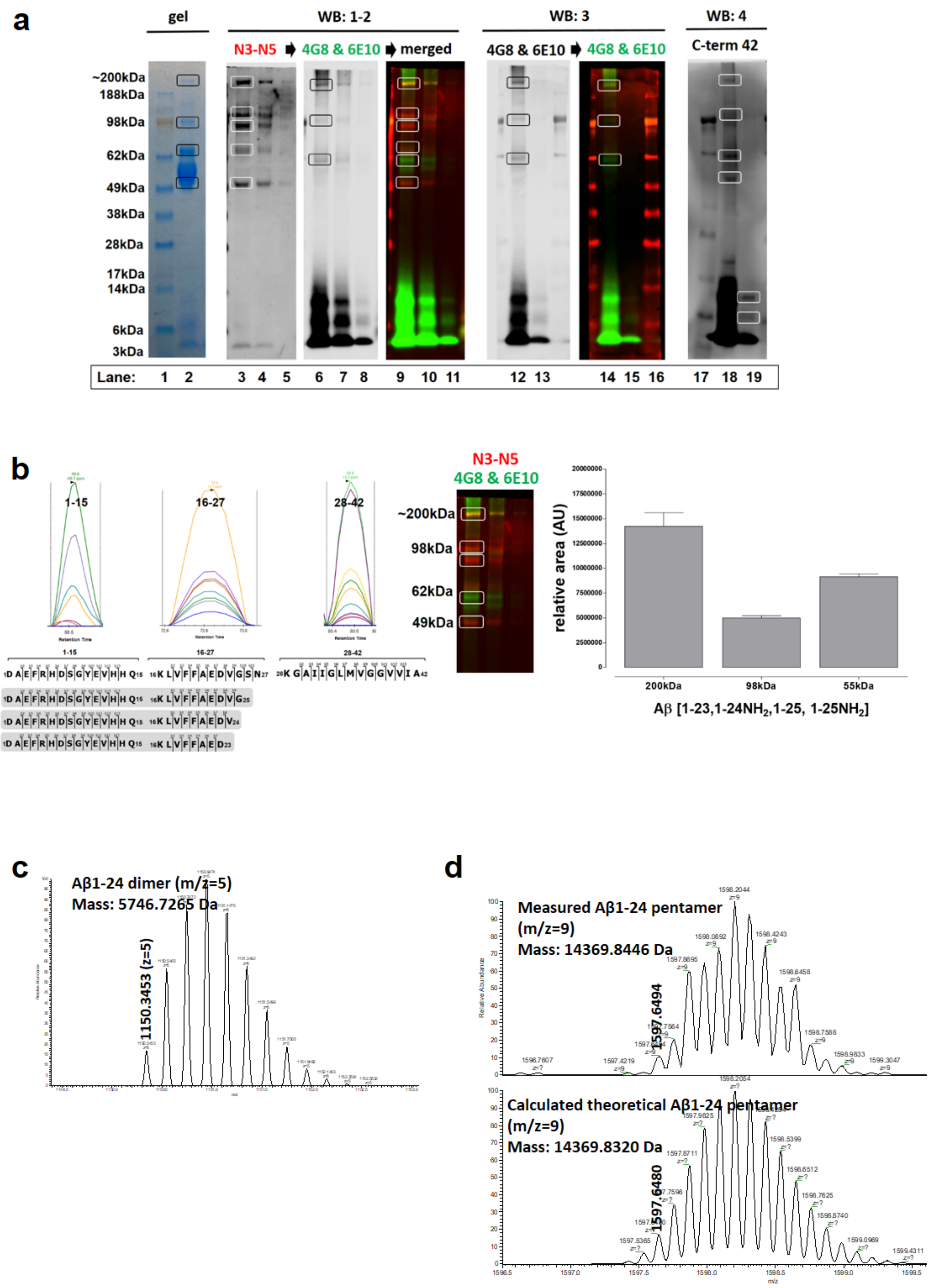


**Supplementary Data Figure 2:**

**a** Same WB as shown in Figure 1c but probed with different antibodies. WB and quantitative MS analysis of high molecular weight (HMW, 50kDa to 200kDa) Aβ assemblies detected in the insoluble fraction. WB were probed using either a combination of 6E10&4G followed by probing with a mixture of neo-epitope antibodies (N3, N4, N5ns & N5NH$_2$) to detect the presence of N-terminal fragments Aβ

1-23 to 1-25 in these HMW oligomers. Gel bands at the migration level of different HMW oligomers were excised and subjected to LysN digestion. (Lanes 1-19 are identical Aβ samples but loaded in triplicates at high to low concentrations. lane: 1, 16, 17 = MWM). (WB: 1-2 was probed sequentially with n3-N5 followed by 6E10&4G8, WBs 3 & 4 were probed individually with 6E10&4G8 and the C-term (Aβ42) (MM26-2) specific antibody respectively). **b** Proteolytic products were subjected to PRM (left) analysis and the relative, calculated peptide areas were used to estimate the relative abundance (right) of the different Aβ fragments (1-23 to 1-25) present in oligomers detected at the gel migration level of ~55kDa, 98kD and ~200kDa. **c** LC-MS measured isotopic distribution of Aβ1-24 dimers and **d** pentamers (top) with the calculated theoretical isotopic distribution (below).

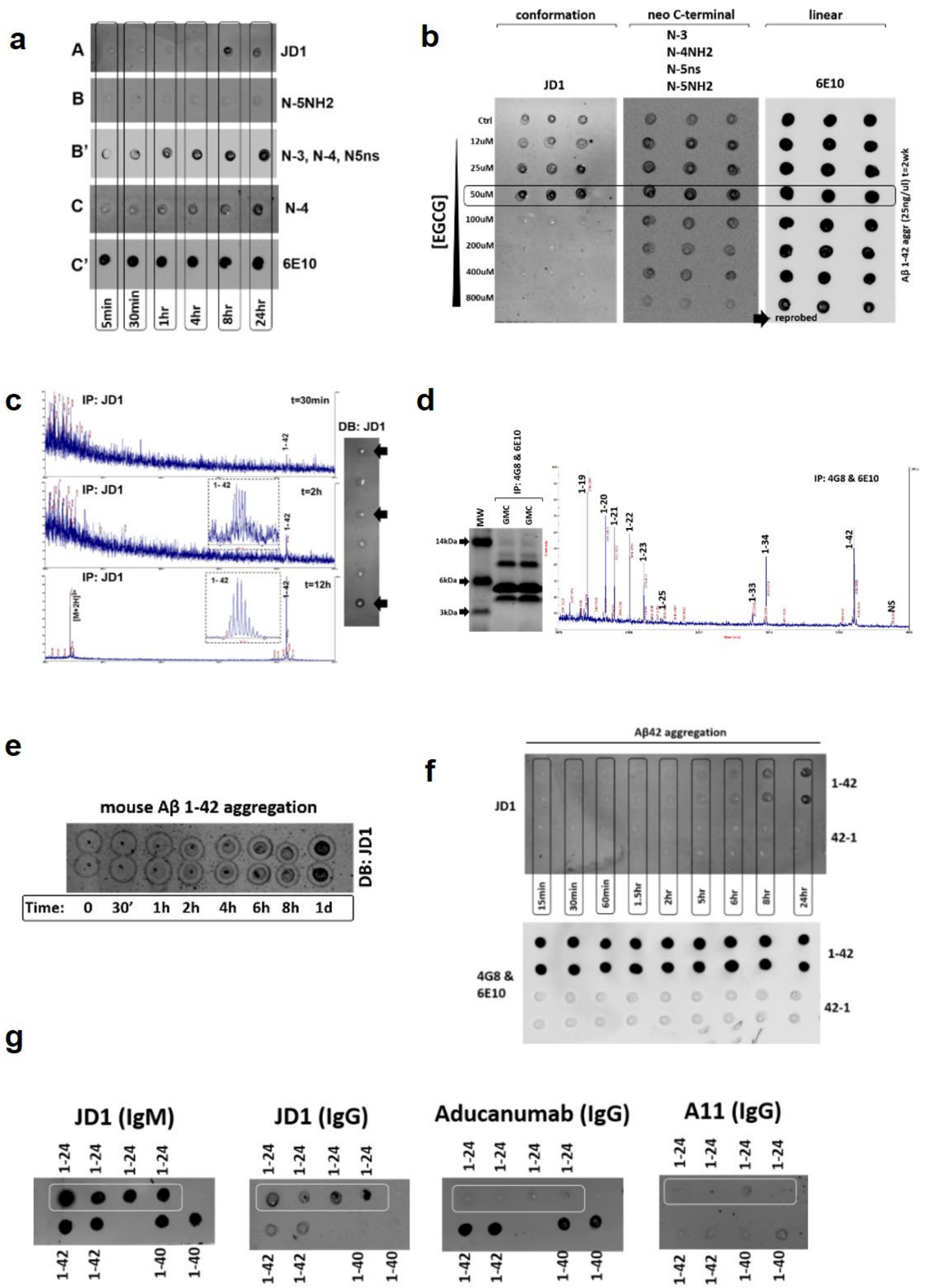


**Supplementary Data Figure 3: a** DB validation and comparison of Aβ42 aggregation kinetics probed with JD1, neo-epitope antibodies (N3, N4, N5ns, N5NH2) and 6E10.

**b** DB screening of Aβ42 aggregation using JD1 in the presence of an aggregation inhibitor (green tea extract EGCG). The degree of inhibition by EGCG can be monitored using both JD1 as well as a combination of neo-epitope antibodies, whereas 6E10 does not detect any time dependent changes in Aβ morphology. The significant decrease in DB signal with ctrl samples (no EGCG: top row) indicates that following 2wks of peptide aggregation, significant change in oligomer morphology is observed and this conversion in morphology can be attenuated with increasing concentrations of EGCG. **c** IP-MS analysis (pull-down) of JD1 positive Aβ assemblies following short-time (30min, top) and prolonged aggregation times (12hrs, bottom). **d** WB and IP-MS analysis of JD1 positive species (Aβ42) found in aged GMC101 worms. The presence of different fragment signatures can be observed, similar to Aβ fragments observed in Tg mouse and human AD brains. **e** DB analysis of aggregated (synthetic) mouse Aβ42 showing the detection of JD1 positive species. **f** DB analysis of Aβ1-42 aggregation kinetics (top) as well as the reversed (below) sequence of Aβ42-1. **g** DB screening of Aβ1-40 & Aβ1-42 and Aβ1-24 oligomers using different conformation specific antibodies as well as both JD1 antibody classes of IgM and IgG. The difference in DB signal intensity reading observed for IgM versus the IgG class of JD1 may account for the pentameric structure and therefore increased secondary antibody binding and hence signal readout. Both IgG and IgM antibody classes show strong binding to fragment Aβ1-24, which was not observed with aducanumab or A11.

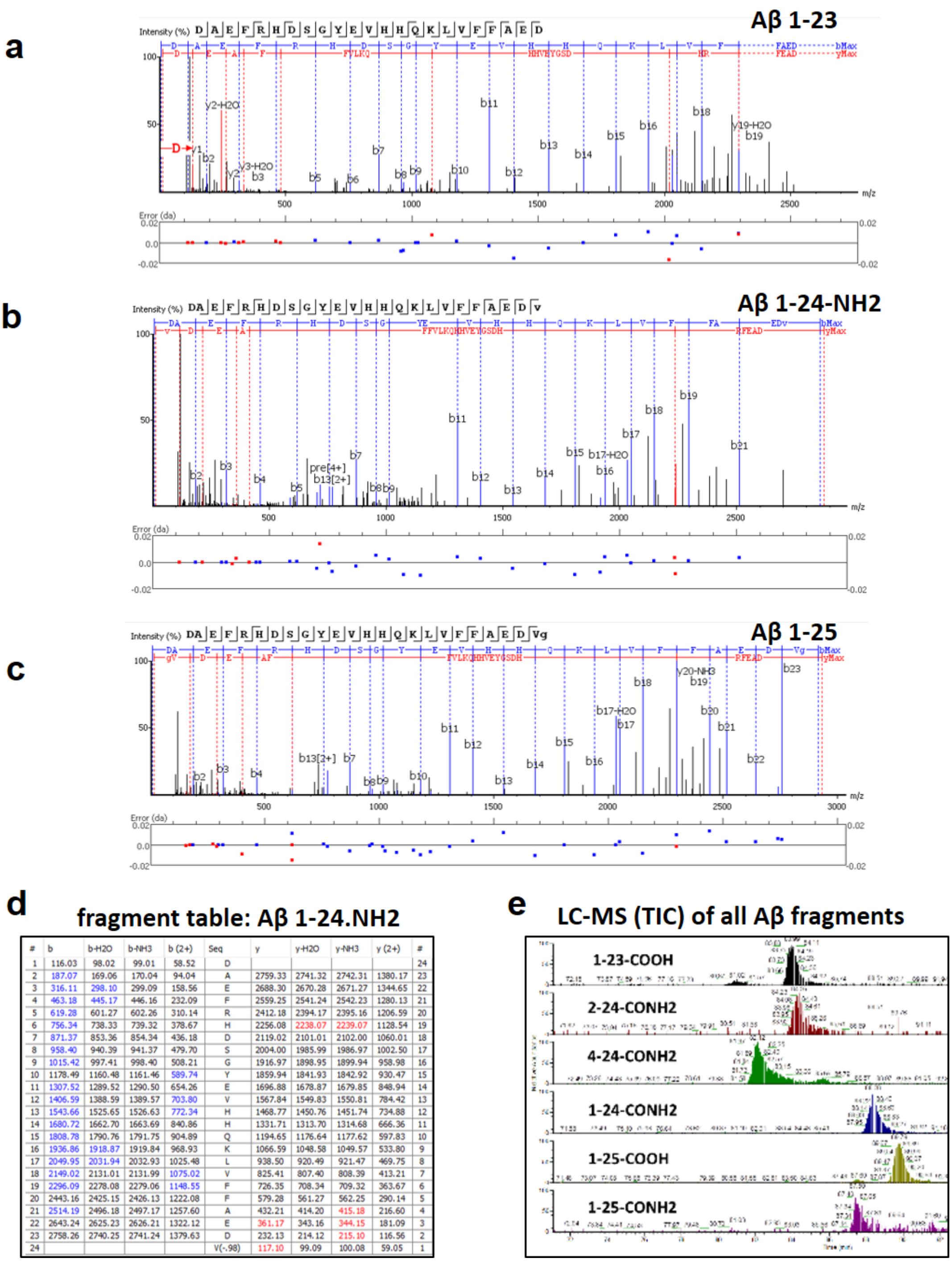


| # | b | b-H2O | b-NH3 | b (2+) | Seq | y | y-H2O | y-NH3 | y (2+) | # |
|---|---|---|---|---|---|---|---|---|---|---|
| 1 | 116.03 | 98.02 | 99.01 | 58.52 | D | | | | | 24 |
| 2 | 187.07 | 169.06 | 170.04 | 94.04 | A | 2759.33 | 2741.32 | 2742.31 | 1380.17 | 23 |
| 3 | 316.11 | 298.10 | 299.09 | 158.56 | E | 2688.30 | 2670.28 | 2671.27 | 1344.65 | 22 |
| 4 | 463.18 | 445.17 | 446.16 | 232.09 | F | 2559.25 | 2541.24 | 2542.23 | 1280.13 | 21 |
| 5 | 619.28 | 601.27 | 602.26 | 310.14 | R | 2412.18 | 2394.17 | 2395.16 | 1206.59 | 20 |
| 6 | 756.34 | 738.33 | 739.32 | 378.67 | H | 2256.08 | 2238.07 | 2239.07 | 1128.54 | 19 |
| 7 | 871.37 | 853.36 | 854.34 | 436.18 | D | 2119.02 | 2101.01 | 2102.00 | 1060.01 | 18 |
| 8 | 958.40 | 940.39 | 941.37 | 479.70 | S | 2004.00 | 1985.99 | 1986.97 | 1002.50 | 17 |
| 9 | 1015.42 | 997.41 | 998.40 | 508.21 | G | 1916.97 | 1898.95 | 1899.94 | 958.98 | 16 |
| 10 | 1178.49 | 1160.48 | 1161.46 | 589.74 | Y | 1859.94 | 1841.93 | 1842.92 | 930.47 | 15 |
| 11 | 1307.52 | 1289.52 | 1290.50 | 654.26 | E | 1696.88 | 1678.87 | 1679.85 | 848.94 | 14 |
| 12 | 1406.59 | 1388.59 | 1389.57 | 703.80 | V | 1567.84 | 1549.83 | 1550.81 | 784.42 | 13 |
| 13 | 1543.66 | 1525.65 | 1526.63 | 772.34 | H | 1468.77 | 1450.76 | 1451.74 | 734.88 | 12 |
| 14 | 1680.72 | 1662.70 | 1663.69 | 840.86 | H | 1331.71 | 1313.70 | 1314.68 | 666.36 | 11 |
| 15 | 1808.78 | 1790.76 | 1791.75 | 904.89 | Q | 1194.65 | 1176.64 | 1177.62 | 597.83 | 10 |
| 16 | 1936.86 | 1918.87 | 1919.84 | 968.93 | K | 1066.59 | 1048.58 | 1049.57 | 533.80 | 9 |
| 17 | 2049.95 | 2031.94 | 2032.93 | 1025.48 | L | 938.50 | 920.49 | 921.47 | 469.75 | 8 |
| 18 | 2149.02 | 2131.01 | 2131.99 | 1075.02 | V | 825.41 | 807.40 | 808.39 | 413.21 | 7 |
| 19 | 2296.09 | 2278.08 | 2279.06 | 1148.55 | F | 726.35 | 708.34 | 709.32 | 363.67 | 6 |
| 20 | 2443.16 | 2425.15 | 2426.13 | 1222.08 | F | 579.28 | 561.27 | 562.25 | 290.14 | 5 |
| 21 | 2514.19 | 2496.18 | 2497.17 | 1257.60 | A | 432.21 | 414.20 | 415.18 | 216.60 | 4 |
| 22 | 2643.24 | 2625.23 | 2626.21 | 1322.12 | E | 361.17 | 343.16 | 344.15 | 181.09 | 3 |
| 23 | 2758.26 | 2740.25 | 2741.24 | 1379.63 | D | 232.13 | 214.12 | 215.10 | 116.56 | 2 |
| 24 | | | | | V(-.98) | 117.10 | 99.09 | 100.08 | 59.05 | 1 |

**Supplementary Data Figure 4: a-c** Summary of MS/MS spectra of identified Aβ fragments Aβ1-23, Aβ1-24 and Aβ1-25 from IP'ed (N3-N4-N5ns) human AD brain samples. **d** MS/MS fragmentation table of theoretical and identified (blue = b-ios and red = y-ions) fragment ions from Aβ1-24. **e** Total ion chromatogram extraction and LC retention times of the different Aβ fragments identified showing N-term truncations at position 2 and 4 as well as C-terminal amidation identified for Aβ1-24 and Aβ1-25.

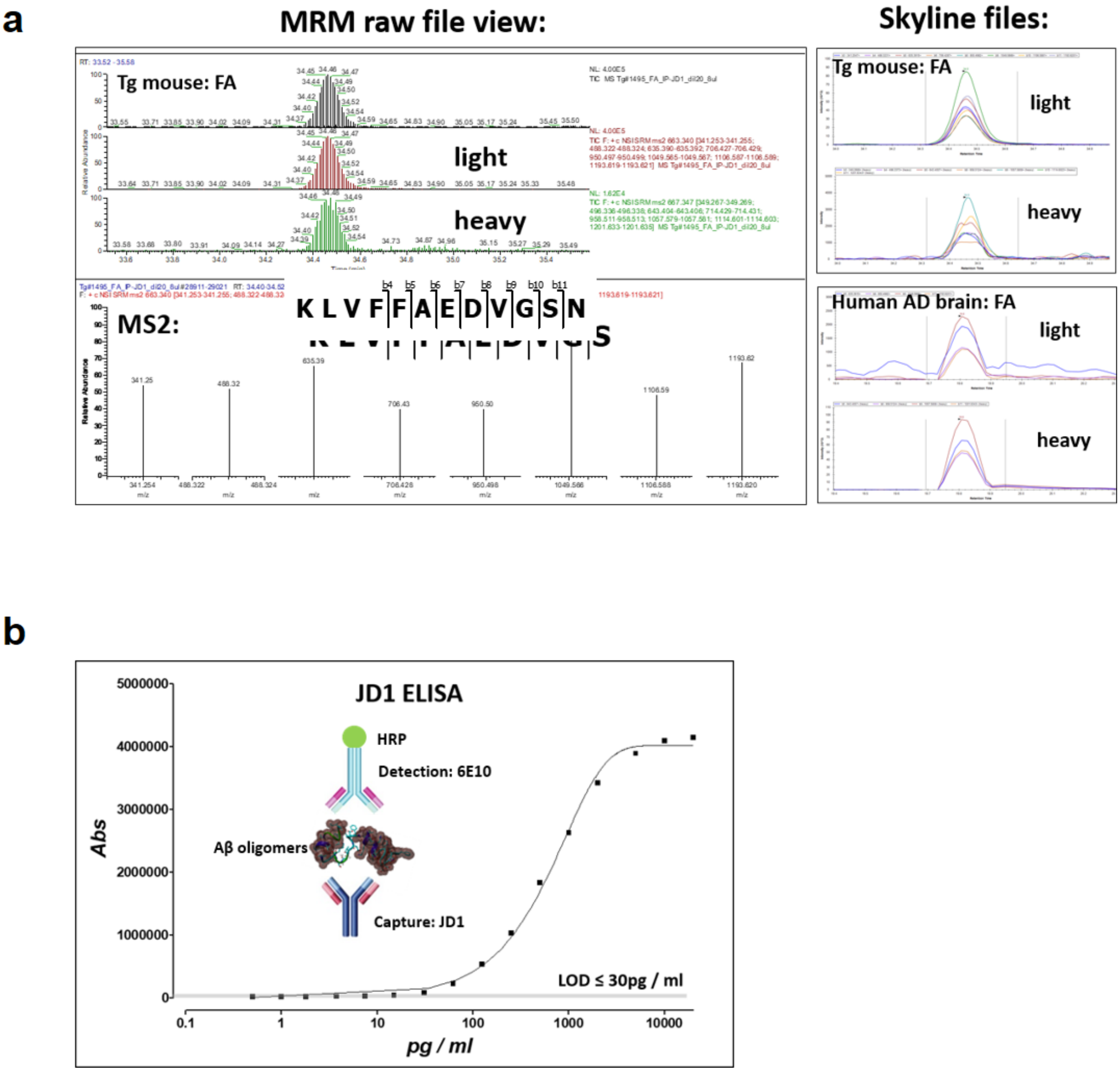


**Supplementary Data Figure 5:**

**a** Quantitative, multiple reaction monitoring (MRM) mass spectrometry raw spectrum of the proteolytic (LysN) cleavage product of fragment Aβ16-27 and right; peak area evaluation using Skyline software. Inset right; example of an IP-MRM (JD1) analysis of Tg2576 mouse (top) and human AD brain (bottom) tissue extracts, showing the calculated peak areas of the endogenous (light) and heavy isotope labelled, (spiked) Aβ fragments. **a** Inset; diagram of the sandwich ELISA concept using JD1 as capture and a biotinylated form of 6E10 as detection antibody as well as a representative titration curve and limit of detection (LOD) of an Aβ1-24 oligomer preparation used for the assay.

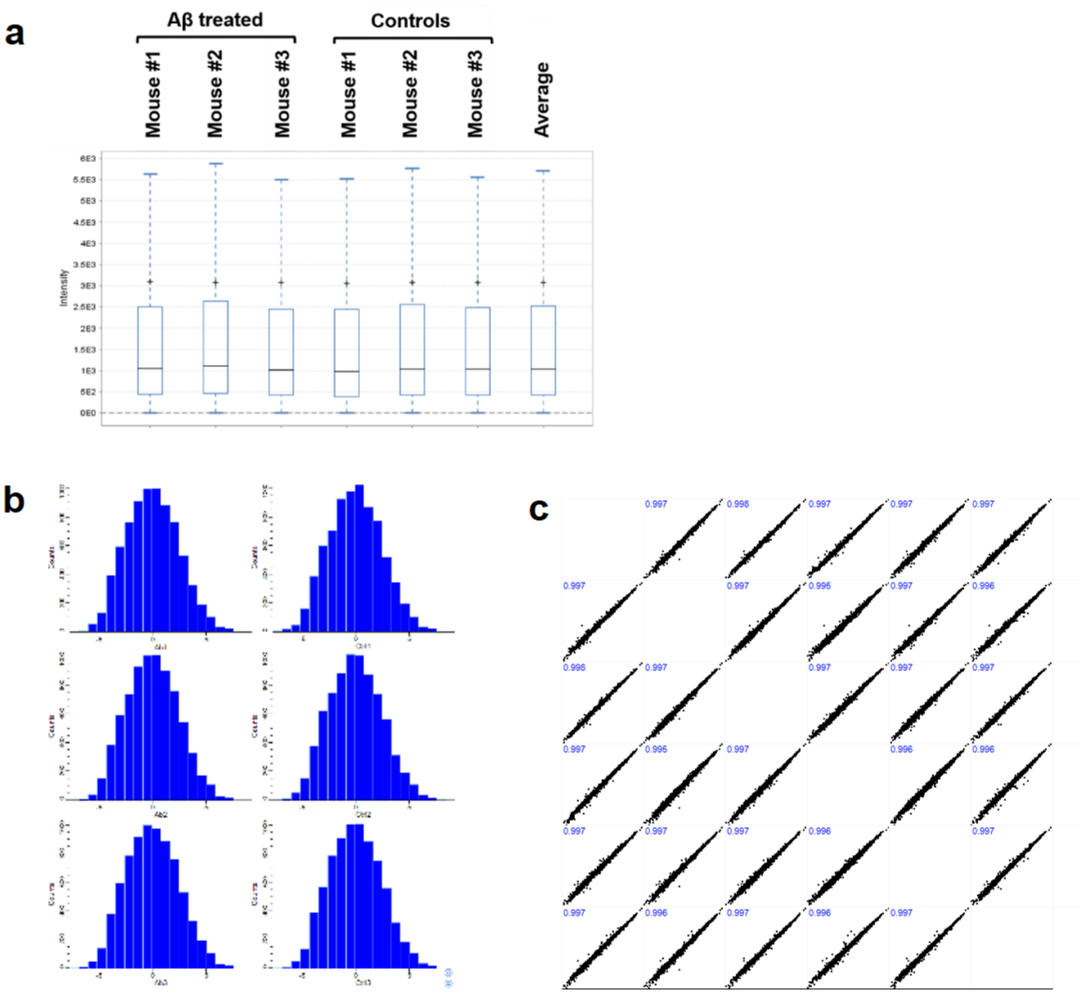


**Supplementary Data Figure 6:**

**a** Summary of overall normalized protein intensity levels following MS signal normalization (PEAKS Studio soft.) showing the median and inter-quantile range of relative MS intensities across all different samples. **b** Protein distribution analysis across all samples (Perseus soft.). **c** Proteome correlation analysis across all analysed samples (PEAKS Studio soft).

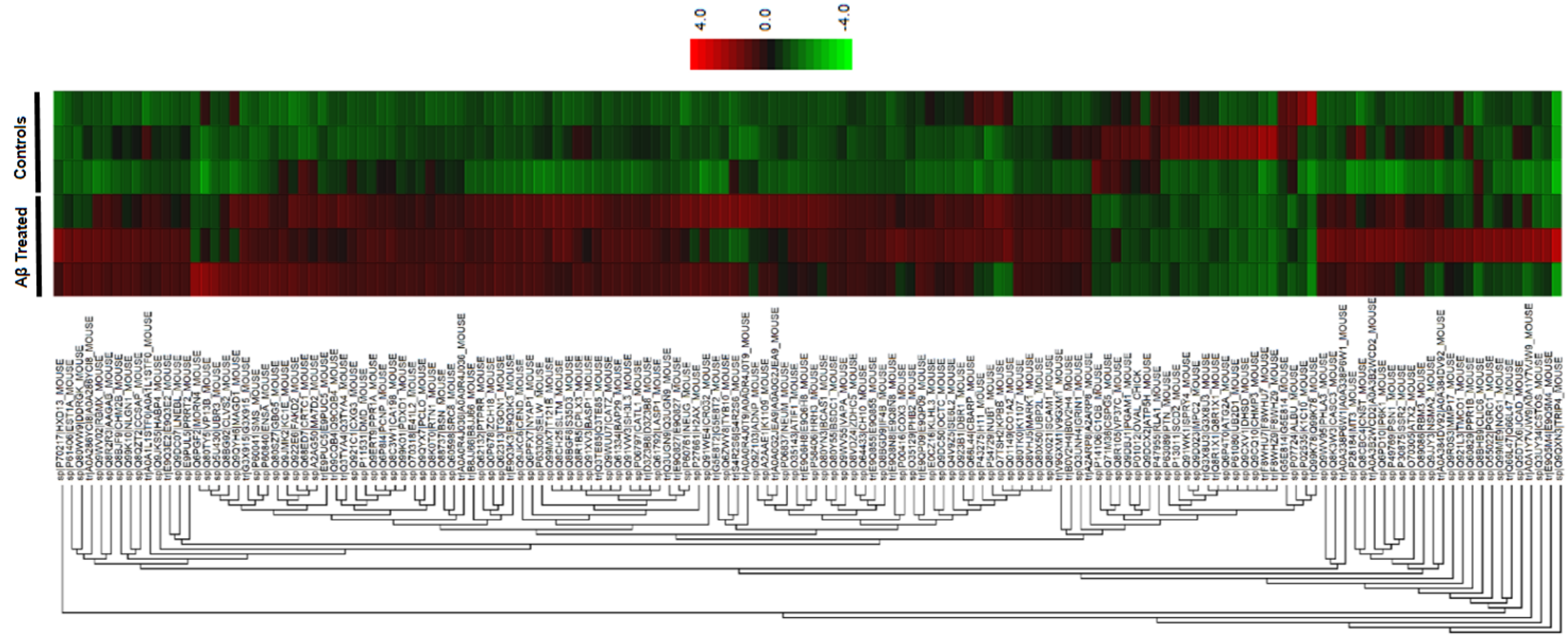


**Supplementary Data Figure 7:** Hierarchical clustering of the relative protein abundance found in mouse brain tissue treated with LMW Aβ oligomers or PBS controls (PEAKS Studio soft.). The hierarchical clustering of proteins is represented as a heat map depicting relative protein abundance (normalized values logged to base 2) of the protein list with filters. The hierarchical clustering is measured with a Euclidean distance similarity measurement of the log2 ratios of the samples relative to a canonical sample.

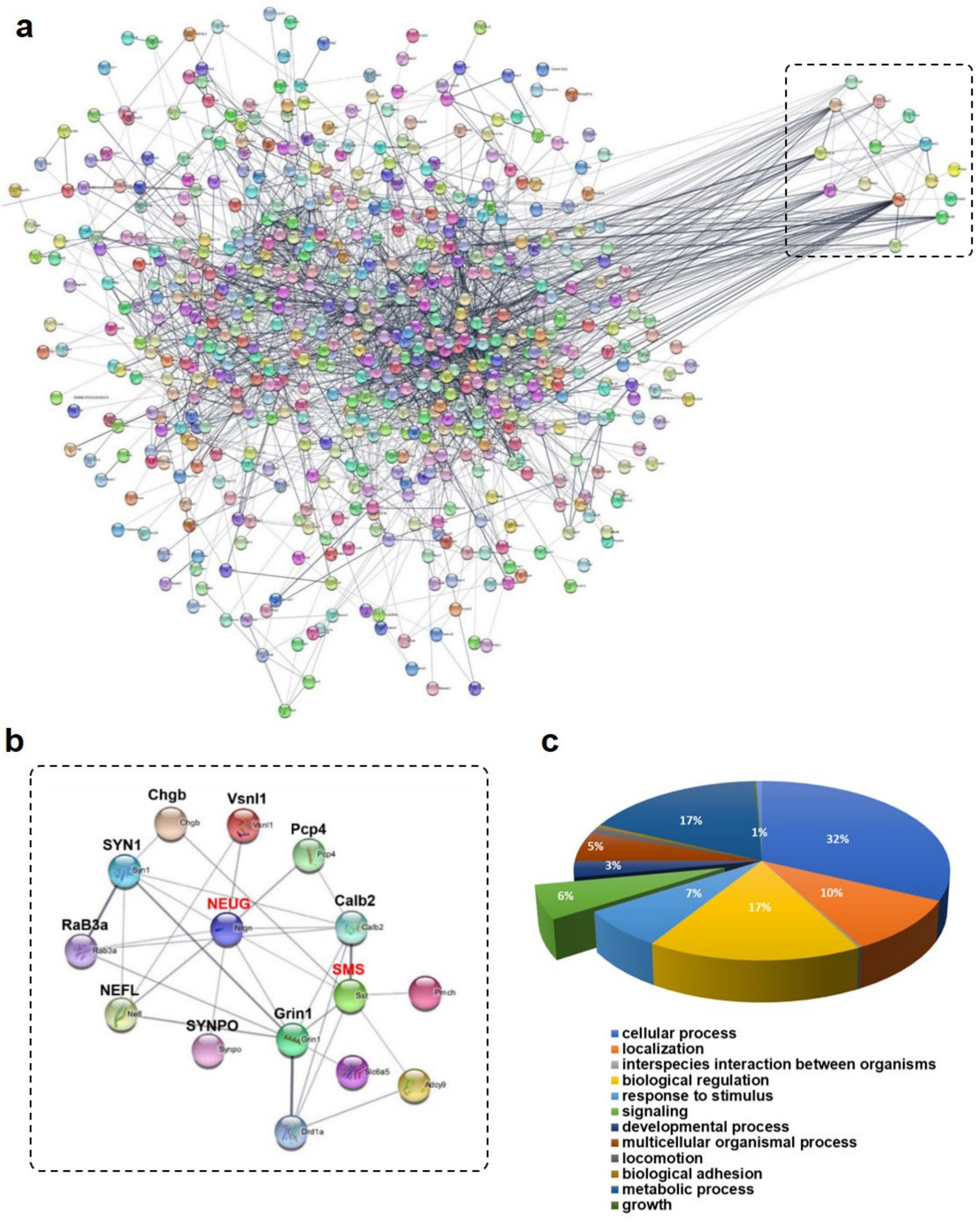


**Supplementary Data Figure 8: a** Protein network analysis (STRING) showing the interaction of >500 proteins identified in both control and Aβ oligomers treated mouse brain tissues. **b** A zoom of the protein association network analysis (STRING), centred on neurogranin (NEUG) protein and showing an interaction with somatostatin (SMS) protein as well as a network with other proteins. **c** Identification of different protein functional groups, of which protein signalling networks accounted for 6%.

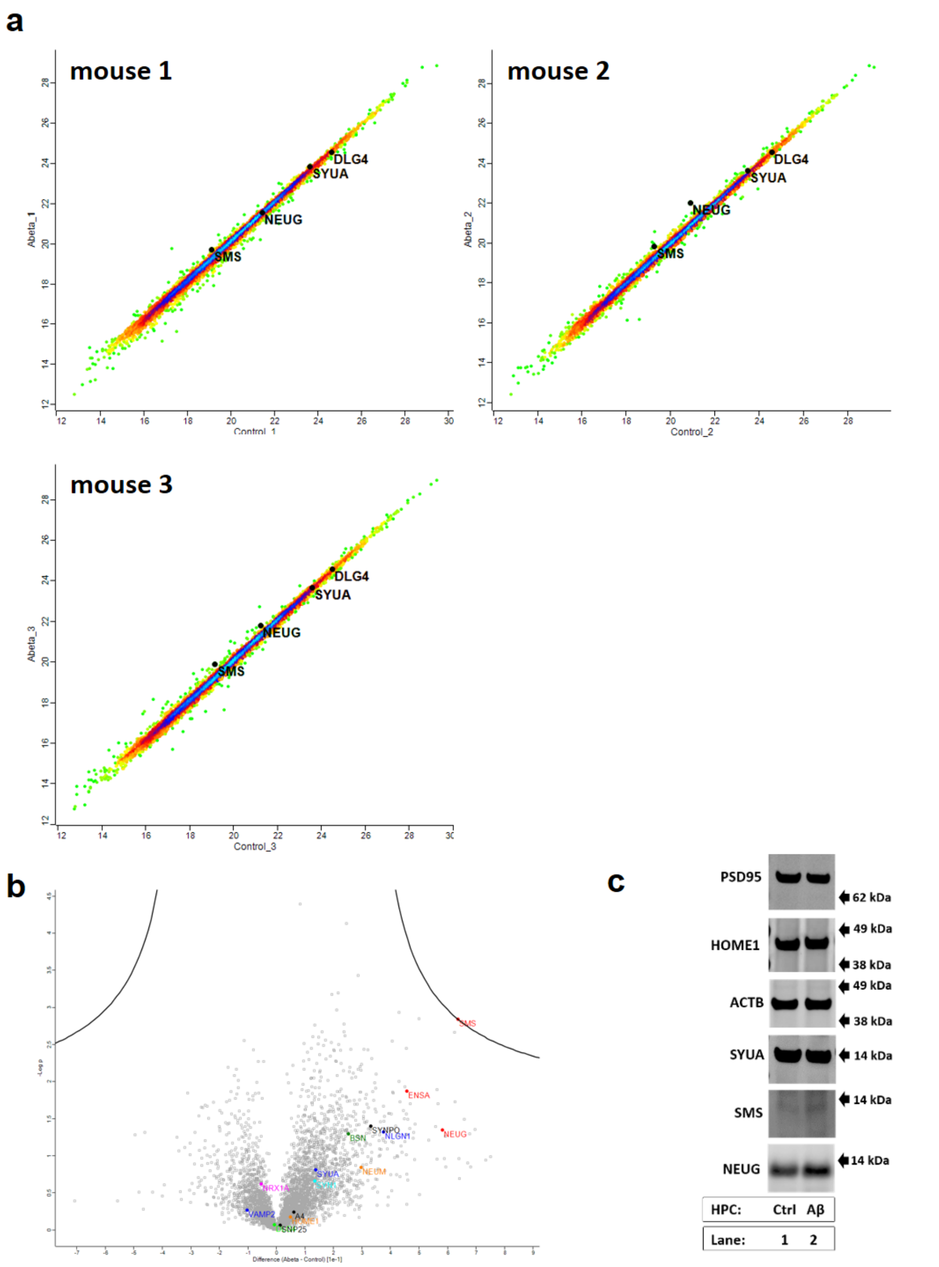


**Supplementary Data Figure 9: a** Intra-TMT channel correlation analysis (density maps) of proteins identified in control (x-axis) vs. Aβ (y-axis) treated mouse brain tissues (left to right: mouse #1 to 3). DLG4 (PSD95) or SYUA proteins do not change significantly and therefore show a strong correlation between control versus Aβ treated animals, whereas MS signal intensity of the two synaptic marker SMS and NEUG respectively appear slightly increased in Aβ treated animals. **b** Volcano plot (zoom), highlighting fold-changes of different pre- and post-synaptic markers. (Perseus soft.). **c** Orthogonal validation of different synaptic proteins using WB analysis, showing the relative abundance of different synaptic markers in a control and Aβ treated mouse brain. No significant changes in levels of synaptic markers were observed between control and Aβ treated tissues except for NEUG (Lane 2). WB detection of SMS proofed to be difficult using two different antibodies (Table I).

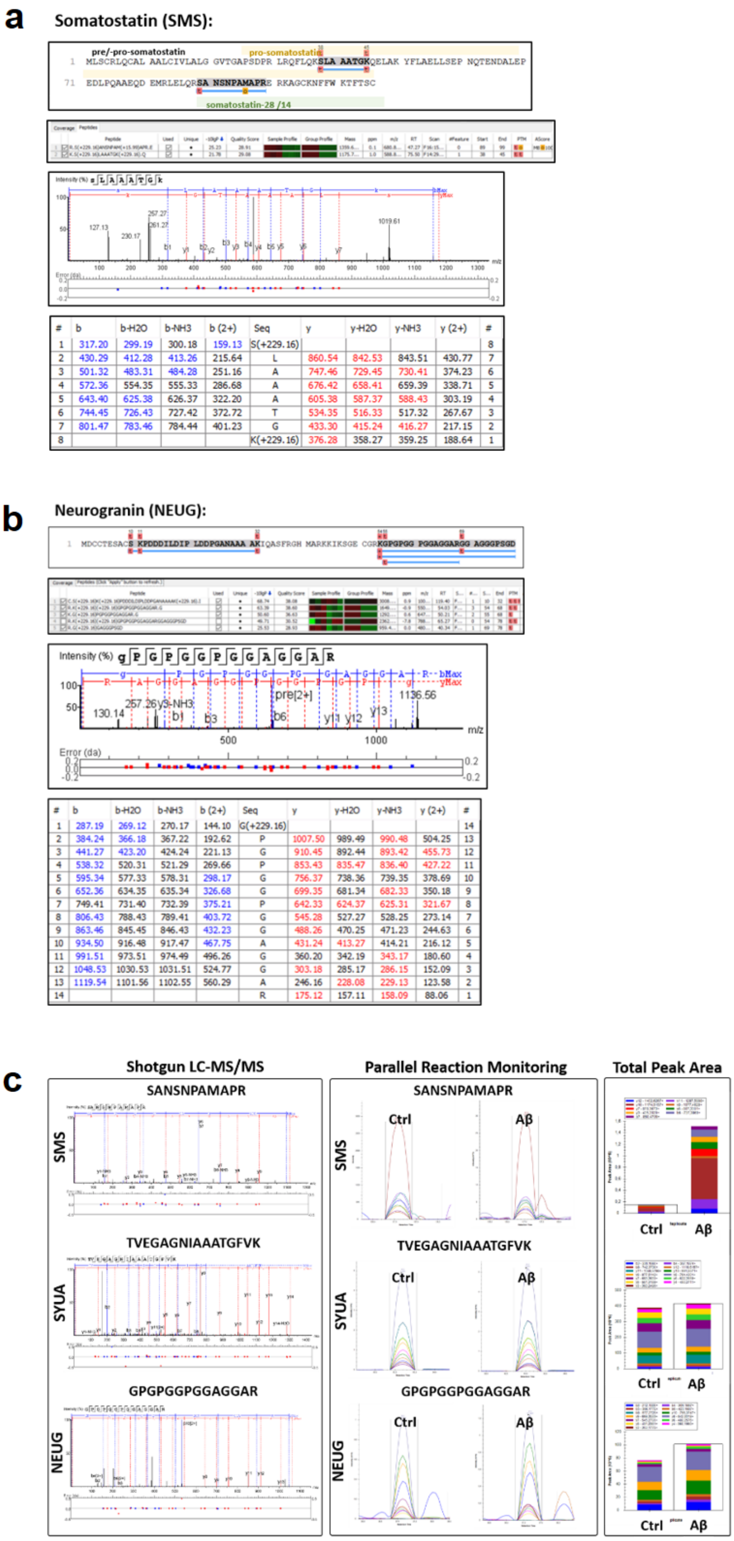


**Supplementary Data Figure 10:** Summary of LC-MS sequencing of two synaptic proteins, showing the overall sequence coverage and identification of two proteotypic peptides for somatostatin (top) **(a)** and below, neurogranin **(b)**.

| No. | Protein ID | Peptide sequence | From | To | Phospho-Site | Ctrl. | Aβ |
|---|---|---|---|---|---|---|---|
| 1 | CAMKIIa | K.E(p)SSESTNTTIEDEDTK.V | 329 | 344 | S330 | | |
| 2 | CAMKIIa | K.ES(p)SESTNTTIEDEDTK.V | 329 | 344 | S331 | | |
| 3 | CAMKIIa | K.ESSESTNT(p)TIEDEDTK.V | 329 | 344 | T337 | | |
| 4 | CAMKIIa | K.AGAYDFP(p)SPEWDTVTPEAK.D | 227 | 245 | S234 | | |
| 5 | CAMKIIa | R.LHD(p)SISEEGHHY.L | 75 | 86 | S78 | | |
| 6 | CAMKIIa | K.ESSES(p)TNTTIEDEDTK.V | 329 | 344 | S334 | | ND |
| 7 | CAMKIIa | R.N(p)SKPVHTTILNPH.I | 406 | 418 | S407 | | ND |
| 8 | CAMKIIa | R.QE(p)TVDCLK.K | 284 | 291 | T286 | | |
| 9 | CAMKIIa | R.FYFENLW(p)SR.N | 397 | 405 | S404 | | |
| 10 | DLG4 | R.GNS(p)SGLGFSIAGGTDNPHIGDDPSIFITK.I | 71 | 98 | S74 | ND | |
| 11 | DLG4 | R.EQLMNSS(p)SLGSGTASLR.S | 409 | 424 | S416 | ND | |
| 12 | DLG4 | K.NTYDVVY(p)SLK.V | 234 | 242 | S241 | | |
| 13 | DLG4 | R.YQDEDTPPLEHS(p)SPAHLPN.Q | 14 | 31 | S26 | | ND |
| 14 | NEUM | K.EGDGSATTDAAPAT(p)SPK.A | 82 | 98 | S96 | | |
| 15 | NFL | K.ESEEEEKKEE(p)SAGEEQVAK.K | 522 | 540 | S532 | | |
| 16 | NFL | K.DEPP(p)SEGEAEEEEK.E | 469 | 482 | S473 | | |
| 17 | NFH | K.AK(p)SPVKEDIKPPAEAK.S | 793 | 808 | S495 | | |
| 18 | NFH | K.SPAAVK(p)SPAEAK.S | 721 | 732 | S727 | | |
| 19 | NFH | K.SPAEAK(p)SPIEVK.S | 757 | 768 | S763 | | |
| 20 | NFH | K.HPTDIRPPEQVK(p)SPAK.E | 847 | 862 | S859 | | |
| 21 | NFH | K.AKPLDVK(p)SPEAQTPVQEEAK.H | 827 | 846 | S834 | | |
| 22 | NFH | K.AK(p)SPVKEDIKPPAE.A | 793 | 806 | S795 | | |
| 23 | NFH | K.KEEVK(p)SPVKEEVK.A | 883 | 895 | S888 | | |
| 24 | NFH | K.EDIKPPAEAK(p)SPEK.A | 799 | 812 | S809 | | |
| 25 | NFH | K.EEVK(p)SPVKEEVK.A | 884 | 895 | S888 | | |
| 26 | NFH | K.SPAEAK(p)SPAEAK.S | 601 | 612 | S607 | | |
| 27 | NFH | K.SPAEAK(p)SPAEVK.S | 667 | 678 | S673 | | |
| 28 | NFH | K.SPAEPK(p)SPATVK.S | 577 | 588 | S583 | | |
| 29 | NFH | K.SPIEVK(p)SPEK.A | 763 | 772 | S769 | | |
| 30 | NFH | K.SPAEPK(p)SPAEPK.S | 559 | 570 | S565 | | |
| 31 | NFH | K.SPAEPK(p)SPAEAK.S | 565 | 576 | S571 | | |
| 32 | SYPH | R.LHQV(p)YFDAPSCVK.G | 77 | 89 | Y81 | | ND |
| 33 | SYN1 | R.PAKPQLAQKP(p)SQDVPPPITAAAGGPPHPQLNK.S | 632 | 663 | S643 | ND | |
| 34 | SYN1 | R.QA(p)SISGPAPTK.A | 566 | 576 | S568 | | |
| 35 | SYN1 | R.ASTAAPVASPAAP(p)SPGSSGGGGFFSSLSNAVK.Q | 54 | 85 | S67 | | |
| 36 | SYN1 | R.PVAGGPGAPPAARPPA(p)SP(p)SPQR.Q | 535 | 556 | S551 / S553 | | |
| 37 | SYN1 | R.PVAGGPGAPPAARPPA(p)SPSPQR.Q | 535 | 556 | S551 | | |
| 38 | SYN1 | P.PQKPPGPAGP(p)TRQ.A | 591 | 603 | T600 | | |
| 39 | SYN1 | R.RL(p)SDSNFMANLPNGYMTDLQR.P | 7 | 27 | S9 | | |
| 40 | SYN2 | R.TPAL(p)SPQRPLTTQQPQSGTLK.E | 422 | 442 | S426 | ND | |
| 41 | TAU | R.HLSNVSSTG(p)SIDMVDSPQLATLADEVSASLAK.Q | 396 | 427 | S405 | | |
| 42 | TAU | K.TDHGAEIVYK(p)SPVVSGDTSPR.H | 375 | 395 | S385 | | |
| 43 | TAU | K.IG(p)SLDNITHVPGGGNK.K | 343 | 358 | S345 | | |
| 44 | TAU | R.HLSNVSS(p)TGSIDMVDSPQLATLADEVSASLAK.Q | 396 | 427 | T403 | | ND |
| 45 | TAU | K.S(p)TPTAEDVTAPLVDER.A | 57 | 72 | T58 | | |
| 46 | TAU | K.TDHGAEIVYK(p)SPVVSGD(p)TSPR.H | 375 | 395 | S385 / T392 | | |
| 47 | TAU | K.TDHGAEIVYK(p)SPVVSGDT(p)SPR.H | 375 | 395 | S385 / S393 | | |
| 48 | TAU | K.VAVVR(p)TPPK(p)SPSASK.S | 215 | 229 | T220 / S224 | ND | |
| 49 | TAU | R.SGYS(p)SPGSPGTPGSR.S | 184 | 198 | S188 | | ND |
| 50 | MARCKS | K.EAAEAEPAEPS(p)SPAAEAEGASASSTSSPK.A | 102 | 130 | S113 | | |
| 51 | MARCKS | K.KE(p)SGEGAEAEGATAEGAK.D | 169 | 186 | S171 | | |
| 52 | MARCKS | K.EAAEAEPAEPSSPAAEAEGASASST(p)SSPK.A | 102 | 130 | S127 | | |
| 53 | MARCKS | K.VNGDA(p)SPAAAEPGAK.E | 41 | 55 | S46 | | |
| 54 | MARCKS | K.GEATAERPGEAAVAS(p)SPSK.A | 12 | 30 | S27 | | |
| 55 | MARCKS | K.AEDGAAP(p)SPSSETPK.K | 131 | 145 | S138 | | |

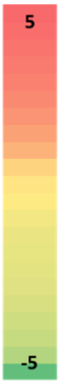


**Supplementary Data Table II:** Summary of MS based identified of site-specific protein phosphorylation in control and Aβ treated mouse brain slices. The intensity heat map represents the relative measured MS1 peak areas of different phosphos-peptides identified in control and Aβ treated brain tissue. Identified phosphosites with increased intensities are marked in red, whereas identifications with lower intensities are marked in green. (ND: not detected)
